# Genomic evidence of early-diverging domesticated lineages in Norwegian farmhouse yeast

**DOI:** 10.64898/2026.03.16.711853

**Authors:** Michael Dondrup, Atle Ove Martinussen, Lisa Karine Haugland, Jonas Brandenburg, Oya Inanli, Abdelhameed Elameen, David Dolan, Sushma Nagaraja Grellscheid, Snorre B. Hagen, Hannes Schroeder, Tor Myking, Hans Geir Eiken

**Affiliations:** Department of Informatics, Computational Biology Unit, University of Bergen, Norway; Western Norway Cultural Academy, Voss, Norway; Division of Biotechnology and Plant Health, Norwegian Institute of Bioeconomy Research, Ås, Norway; Michael Sars Centre, University of Bergen, Norway; Globe Institute, Faculty of Health and Medical Sciences, University of Copenhagen, Denmark; BioArch, Department of Archaeology, University of York, United Kingdom; Division of Environment and Natural Resources, Norwegian Institute of Bioeconomy Research, Ås, Norway; Division of Forestry and Forest Resources, Norwegian Institute of Bioeconomy Research, Ås, Norway; Department of Biosciences, Durham University, United Kingdom

**Keywords:** Kveik, *Saccharomyces cerevisiae*, domestication, yeast evolution, phylogenetics, admixture, population genomics, traditional brewing, structural variation

## Abstract

The use of *Saccharomyces cerevisiae* to ferment alcoholic beverages is an ancient tradition, with genetic evidence indicating origins in Neolithic Asia, although the domestication process of the species is not fully understood. Kveik is a group of traditional yeasts used in farmhouse brewing in western Norway preserved through generations of rural brewing practice. While recent studies have highlighted the distinctiveness of kveik, its precise phylogenetic position, genetic diversity, and domestication history remain unclear. We performed whole-genome sequencing on 64 isolates from 25 traditional brewing cultures from Western Norway selected using cultural heritage criteria, and generated telomere-to-telomere (T2T) assemblies for selected strains. Phylogenomic and population genetic analyses reveal that kveik strains form a paraphyletic and early diverging group with respect to other domesticated *S. cerevisiae* strains. Most strains exhibit low diversity among isolates from the same brewing culture, strong geographic clustering, and little evidence of gene-flow or admixture. Mitochondrial genomes and Ty1 retrotransposon profiles corroborate this distinct lineage history. We further show that previously reported signals of gene flow between kveik and Asian fermentation strains are likely artifacts caused by population structure and selection. Divergence time estimates suggest that the common ancestor of beer, kveik, and other liquid-phase fermenting strains originated from ancestral populations 4,000 to 8,000 years ago. Our genomic resource sheds light on yeast evolution and domestication. Kveik likely comprises some of the oldest domesticated lineages in continuous use until today, connecting intangible cultural heritage to an early genetic origin.

**Significance statement:** Yeast has been used to brew beer and ferment food for thousands of years, but we still do not fully understand how and where the different domesticated strains we are using today arose. We sequenced the largest collection of Norwegian farmhouse yeasts (kveik) to date, from cultures used by rural traditional brewers for generations. Our genetic analyses show that kveik forms a distinct and early branch among European brewing yeasts. This shows that historical traditions have created unique local types of yeast and helps us better understand and preserve the diversity of domesticated yeasts.

## Introduction

The budding yeast *Saccharomyces cerevisiae* has been utilized by humans in the fermentation of various food and beverage products for millennia. Recent phylogenetic evidence suggests that East Asia, particularly China, may represent the center of origin for one or more domestication events (Bai *et al*., 2022). Genomic analyses have revealed that wild and domesticated populations of *S. cerevisiae* form genetically distinct groups. Moreover, domesticated strains can be divided into two major lineages associated with liquid and solid-state fermentations, both of which appear to have originated from a common ancestral population (Duan *et al*., 2018; Han *et al*., 2004; Peter *et al*., 2018).

Today, there are over 2500 sequenced genomes of wild and domesticated isolates of *S. cerevisiae* (Duan *et al*., 2018; Libkind *et al*., 2020; O’Donnell *et al*., 2023). This wealth of data can aid developing new fermentation processes and applications, as well as further understanding this yeast’s evolutionary history and domestication (Libkind *et al*., 2020). Mitochondrial DNA (mtDNA) is also widely used to investigate phylogenetic relationships, and the unusually large mitochondrial genome of S. *cerevisiae* of 75-85 kb may be important to understand its evolution (Wolters *et al*., 2015; Wolters *et al*., 2023). In addition, transposable elements (TEs) are nearly ubiquitous mobile genetic elements that can affect host genome fitness and reflect evolutionary history (Pritham, 2009; Wang *et al*., 2024; Wells & Feschotte, 2020). The compact *S. cerevisiae* genome contains a comparatively small fraction of TEs (<3.3%) belonging to seven known families of long terminal repeat (LTR) retrotransposons all of which have been intensively studied (Bleykasten-Grosshans *et al*., 2013; Carr *et al*., 2012; Czaja *et al*., 2020). Phylogenetic analysis of the Ty1 family of retrotransposons in *S. cerevisiae* may help to resolve biogeography, with the non-canonical Ty1’ element associated with the ancestral wild lineages and the canonical Ty1 element found only in domesticated strains (Czaja *et al*., 2020).

Present industrial brewing yeasts likely share a common ancestor from around 1600-1700 AD, predating modern microbiology (Gallone *et al*., 2018; Gallone *et al*., 2016), but long after the estimated invention of beer brewing around 3000 BC (Gallone *et al*., 2016; McGovern *et al*., 2004; Michel et al. 1992). Phylogenetic analyses divide industrial beer yeast into two major clades: Beer 1, which exhibits the strongest signs of domestication and geographical boundaries, and Beer 2 with more limited traits of domestication (Gallone *et al*., 2016).

Historical sources indicate that farmhouse brewing in Norway has been a living tradition for over a millennium, documented by the oldest laws in Norway (The Gulating Law: Simensen (2021)) and continuing with the present-day tradition of brewing and maintaining local cultures in western Norway (Garshol, 2023; Kits & Garshol, 2021). Western Norwegian yeasts, known as “kveik”, used in farmhouse brewing were typically cultivated and passed down through generations within the same family. In other Nordic countries, such as Finland, the custom of maintaining local cultures is however no longer practiced and the genetic diversity of local yeasts has been lost (Catallo *et al*., 2020). Kveik yeasts have gained global interest due to their unique properties such as fast fermentation, strong flocculation, high temperature and ethanol tolerance, and the production of multiple flavor metabolites with little off-flavors at high temperatures (Foster *et al*., 2022; Kawa-Rygielska *et al*., 2022; Kits & Garshol, 2021; Preiss *et al*., 2018).

Phylogenetic analysis of six Norwegian kveik isolates (Preiss *et al*., 2018) first suggested that these form a genetically distinct group of domesticated *S. cerevisiae* related to the Beer 1 clade, but with conserved mixed ancestry that is not observed within any other beer yeast group. Recently, kveik yeasts were proposed to belong to a larger group of European farmhouse yeasts with Asian admixture, forming a sister group to Beer 1 (Preiss *et al*., 2024). Previous studies on kveik yeasts also suggested mixtures of different *S. cerevisiae* strains in kveik brewing cultures (Preiss *et al*., 2018), hybridization with other yeast species (Krogerus *et al*., 2018), and Asian admixture (Preiss *et al*., 2024). Thus, we intended to test whether our kveik yeast cultures were homogeneous, if there were any recent genetic hybridizations or introgressions, and if there existed traces of novel or ancestral admixture in the kveik yeast.

To resolve the uncertainty of the origin of kveik, we collected putative kveik yeast from traditional brewers in western Norway guided by cultural heritage criteria. We performed whole genome sequencing on isolates from single yeast colonies from which four of the samples were also sequenced using long-read sequencing by Oxford Nanopore technology (ONT) followed by *de novo* genome assembly, variant detection, phylogenetics and population genomics analyses.

## Results

### The variation landscape of kveik-yeast strains

We collected 25 putative kveik yeasts original brewing cultures from traditional brewers in western Norway (Figure 1, Table 1) guided by cultural heritage criteria (see Materials and Methods). From these we isolated 64 single yeast colonies and performed whole genome sequencing using Illumina short-read sequencing where four of these isolates also were sequenced using long-read Oxford Nanopore technology. We included a panel of selected isolates from the panel compiled by Tellini *et al*. (2024) resulting in a total of 302 samples. After *de novo* variant calling and quality filtering, we retained 136,962 high-quality SNPs and 5,313 indels (Additional Data VCF Files). Norwegian and Baltic isolates consistently showed the highest heterozygosity of all genomes, with a heterozygosity-ratio in [0.28, 0.48] (sample 7-1b and sample 4-1a, respectively) whereas all other strains ranged in [0, 0.3] (ERR1308813 - Wine and ERR1308839 – Beer, respectively) (Supplementary Data 5).

**Figure 1.**
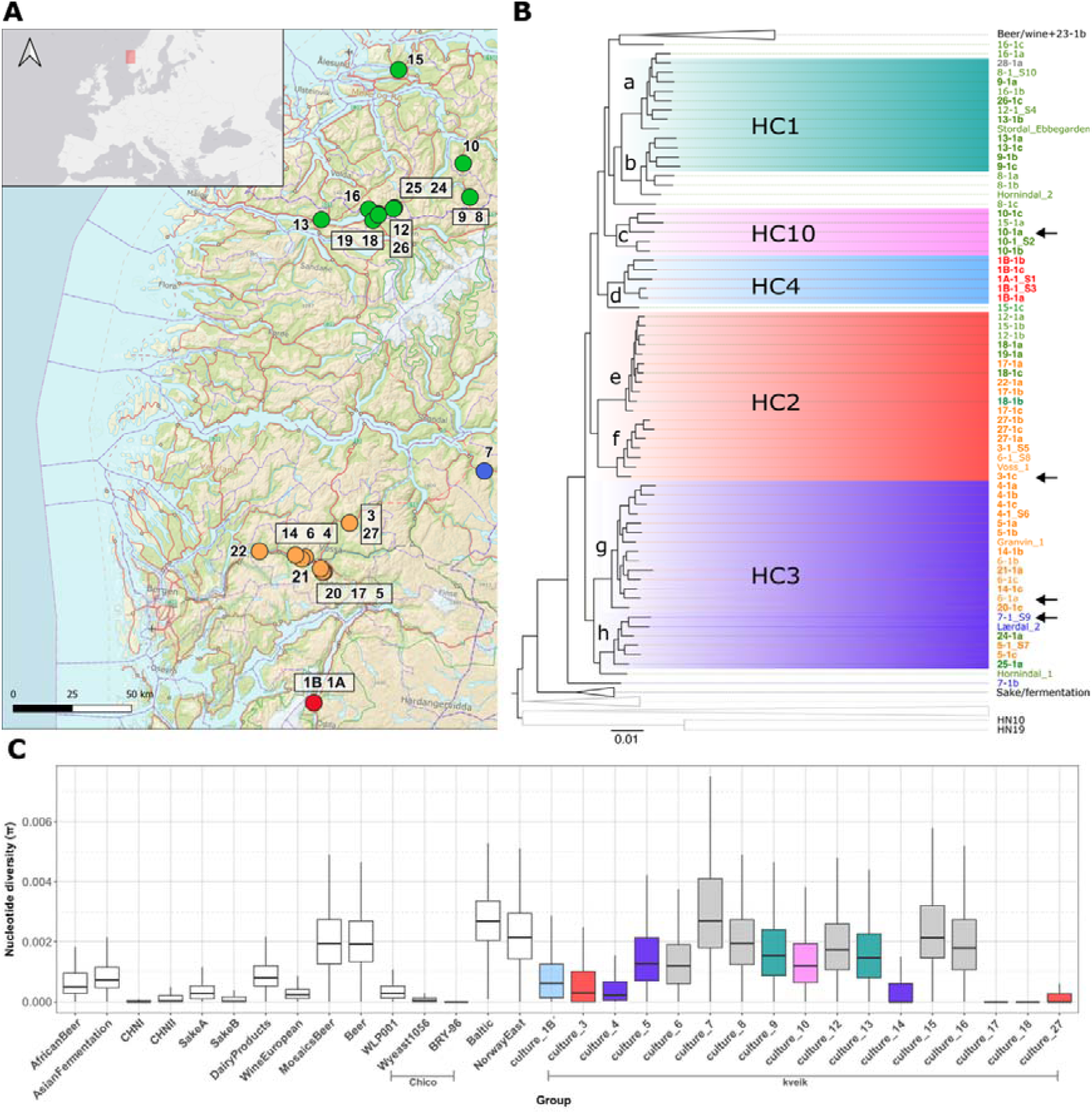
Geographic origin of the Western Norwegian kveik yeast traditional brewing cultures (A), focused maximum-likelihood phylogenetic analysis of 62 isolated kveik yeasts based on 287,696 parsimony-informative sites (B) and boxplot of within group nucleotide diversity across the kveik yeast genomes together with data from previously published genome sequences (C). The full phylogenetic tree is given in Supplementary Figure 1; the full list of included taxa is given in Supplementary Data 9. Colors of taxon labels (orange, green, blue and red) represent the major geographic regions of sampling (see Table 1). Sample IDs of our samples correspond to the “Sample ID” column in Table 1. The identifier of the individual isolate is appended to the numerical ID (e.g. 27-1a, 27-1b, etc.). The sampling locations of the original brewing cultures are indicated by circles (A), and taxon labels represent isolates (B). Compact subclades within the kveik group were visually identified and labeled a-h in the tree, and the resulting genetic kveik cluster designations are as follows: homogeneous cluster (HC) 1 (a & b), HC10 (c), HC4 (d), HC3 (g & h) and HC2 (e & f). Taxon names of samples that have been unanimously assigned to a HC are printed in bold. Isolates that have been sequenced by nanopore technology are indicated by arrows. Nucleotide diversities are calculated in overlapping 10 kilobase windows across the genome for all kveik brewing cultures having two or more isolates and selected groups for comparison. Selected groups contain 12 previously published yeast stains (see Gallone et al. (2016), Preiss et al. (2018), and Duan et al. (2018)) and three commercial monocultures with multiple sequences related to the ‘Chico’ group of ale yeasts (WLP001, Wyeast1056, BRY-96) (C). Boxes of kveik groups are colored according to HC (or grey if isolates from the same culture fell into different clusters) Basemap (A): © Norwegian Mapping Authority / Geonorge (WMS: Topographic Norway Map, permission granted) Source: https://wms.geonorge.no/skwms1/wms.topo

**Table 1.** Kveik yeast samples from traditional brewing cultures used in the study. Sample ID colors refer to positions in map and phylogenetic tree (see Figure 1). Column n gives the number of different isolates sequenced from each brewing culture. Nucleotide diversity was calculated only for brewing cultures having 2 or more sequenced isolates. n.d.= not determined.

| Culture ID | Location | Trivial name | Form | brewing | | since | Diversity ($\pi$) | Note |
| --- | --- | --- | --- | --- | --- | --- | --- | --- |
| 1A | Eitheim, Odda | Eitheim-kveik | Liquid | 2020 | 1 | from grandfather | n.d. |  |
| 1B | Eitheim, Odda | Eitheim-kveik | Dry | 2018 | 4 | from grandfather | 0.00207 | Same origin as 1A |

*Table 1. Kveik yeast samples from traditional brewing cultures used in the study.
|  |  |  |  |  |  |  |  |  |
| --- | --- | --- | --- | --- | --- | --- | --- | --- |
| 3 | Selland, Vossestrand | Selland-kveik 1 | Dry | 2020 | 2 | from grandfather | 0.00120 |  |
| 4 | Røte, Dyrvedalen, Voss | Stolpeklattrar-kveik | Liquid | 2020 | 4 | 1920 | 0.00084 |  |
| 5 | Herre, Bordalen, Voss | Herre-kveik | Liquid | 2020 | 4 | 1990 | 0.00316 |  |
| 6 | Gjernes, Voss | Gjernes-kveik | Liquid | 2020 | 4 | from father | 0.00303 |  |
| 7 | Ljosne, Lærdal | Lærdal-kveik | Dry | 2021 | 2 | 1994 | 0.00757 |  |
| 8 | Geiranger | Geiranger-1 | Dry | 2020 | 4 | 1960 | 0.00510 |  |
| 9 | Geiranger | Geiranger-2 | Dry | 2019 | 3 | 1960 | 0.00443 |  |
| 10 | Hjelle, Eidsdal | Tore Hjelle-kveik | Dry | 2021 | 4 | 1982 | 0.00316 |  |
| 12 | Grodås | Lødemel-kveik | Dry | 2021 | 3 | 1969 | 0.00469 |  |
| 13 | Nordfjordeid | Nordfjordeid-kveik | Dry | 2021 | 3 | 1944 | 0.00427 |  |
| 14 | Hovda, Vestbygdi, Voss | Hovda-kveik | Liquid | 2021 | 2 | before 1987 | 0.00059 |  |
| 15 | Stavset, Ålesund | Pål-kveik | Liquid | 2021 | 3 | from grandfather | 0.00567 |  |
| 16 | Otterdal, Hornindal | Otterdal-kveik | Dry | 2019 | 3 | 1980 | 0.00497 |  |
| 17 | Gjelland, Bordalen, Voss | Gjelland-kveik 1 | Dry | 1991 | 3 | from father | 0.00009 |  |
| 18 | Kjos, Hornindal | Kongsvik-kveik 1 | Dry | n.d., in a box | 3 | from grandfather | 0.00010 |  |
| 19 | Kjos, Hornindal | Kongsvik-kveik 2 | Dry | n.d., in envelope | 1 | from grandfather | n.d. |  |
| 20 | Gjelland, Bordalen, Voss | Gjelland-kveik 2 | Liquid | 2021 | 1 | 2000 | n.d. |  |
| 21 | Græe, Bordalen, Voss | Bordalen-kveik | Dry | 2022 | 1 | n.d. | n.d. |  |
| 22 | Bolstad | Bolstad-kveik | Dry | 2022 | 1 | from great grandfather | n.d. |  |
| 23 | Skaun | Skaun gjær (yeast) | Liquid | 2022 | 1 | n.d. | n.d. | not kveik |
| 24 | Nygård, Grodås | Nygård-kveik | Dry | 2022 | 1 | from father | n.d. |  |
| 25 | Seljeset, Grodås | Stalljen-kveik | Dry | 2022 | 1 | from father in 1970-ties | n.d. |  |
| 26 | Toppen, Grodås | Nesdalkveik | Dry | 2020 | 1 | 1995 | n.d. |  |
| 27 | Selland, Vossestrand | Selland-kveik 2 | Dry | n.d. (est. 50-100 years) | 3 | Same farm as No. 3 | 0.00045 | Historic sample from old barrel |
| 28 | Unkown | Gjeving-kveik | Liquid | 2022 | 1 | n.d. | n.d. | Highly similar to sample 16 |

Among the 64 isolates, 62 were determined to belong to both the phylogenetic and geographical group of kveik yeast (see Figure 1). We estimated nucleotide diversity (π) and Watterson’s theta (θ) across all kveik isolates (n = 62). The estimated nucleotide diversity (π = 0.00586) was higher than Watterson’s theta (θ = 0.00392), resulting in a strongly positive Tajima’s D (D = 1.752). This indicates an excess of intermediate-frequency variants in the combined kveik dataset. Such a signal may indicate balancing selection acting on certain loci, or more specifically, a pronounced population structure, where genetically distinct subgroups (e.g., based on geography or brewing tradition) contribute divergent alleles to the pooled analysis. This prompted us to investigate the underlying population structure of kveik more deeply by attempting to divide it into homogeneous populations.

### Kveik yeast brewing cultures show internal diversity

Previous work indicated that kveik yeast cultures have diverse colony morphology which may reflect internal diversity (Preiss *et al*., 2018). To investigate this, we isolated multiple single colonies as biological replicates from several samples, including two samples (IDs 3 and 27; Table 1) collected at two different time-points, with intermediate reuse in brewing. Notably, we did not observe any deviations in colony morphology and only minor genetic divergence of isolates from the same culture sampled years apart, even after multiple brewing cycles. To assess diversity, we sequenced up to four isolates from 17 brewing cultures either from distinct colonies or independent sampling.. The majority of the original brewing cultures (11 of 17) had isolates that consistently grouped into the same clusters, indicating limited genetic variation and stability within the original brewing culture, while the remaining six original brewing cultures showed higher genetic variation although all within the kveik phylogenetic group (Figure 1B, 2, Supplementary Figure 1). Between-isolate nucleotide diversity π (Figure 1C) was itself variable between cultures indicating different demographic histories, population sizes and bottleneck strengths. We included three repeatedly sequenced commercial mono-cultures related to the ‘Chico’ group (WLP001, Wyeast1056, and BRY-96) as controls. When compared to other larger groups, π was generally lower in kveik than in the total Beer and MosaicsBeer groups (except for isolates 7, 8, 15, and 16) but higher than in commercial mono-cultures, whereas five isolates (4, 14, 17, 18, and 27) showed reduced diversity.

**Figure 2.**
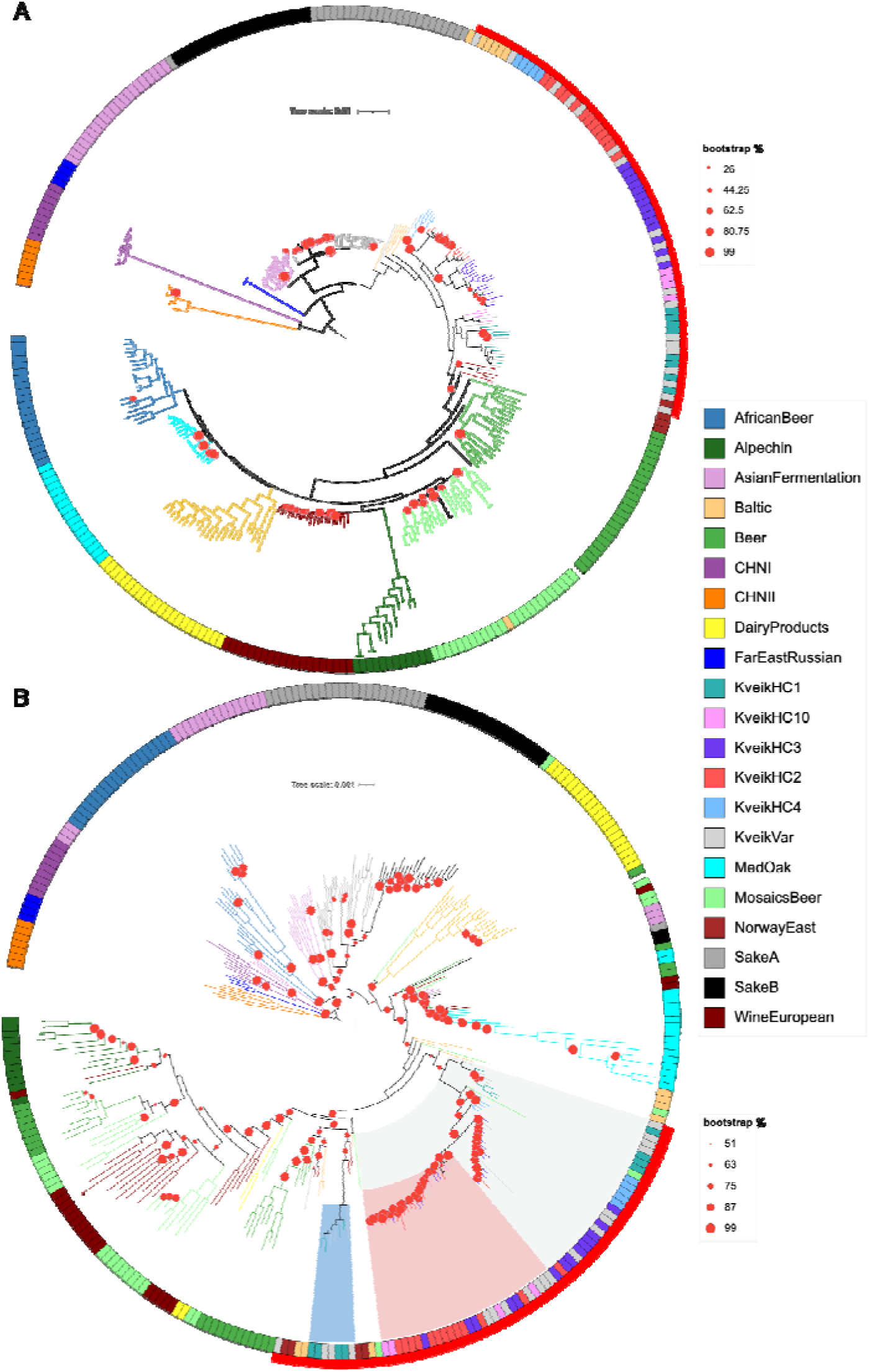
Maximum likelihood phylogenies of the nuclear genomes based on 468,897 parsimony-informative sites under the TVMe+ASC+R8 model (A) and mtDNA based on 5095 parsimony-informative sites under the TVM+F+I+R7 model (B) of 302 S. cerevisiae genomes. Both trees are rooted on the clade containing CHNI/II. All clades except CHNI/II and FarEastRussian are considered domesticated. Nodes with bootstrap support < 100% are marked with red dots of proportional diameter, nodes with 100% support are unmarked. Color codes represent strain groups and are identical in both plots, with the red outer segment marking the extension of kveik yeast. In panel B, clades consisting exclusively of kveik strains are highlighted in grey and blue. A clade of kveik strains with markedly low internal branch length is highlighted in red.

### Telomere-to-telomere and haplotype resolved assemblies

Oxford Nanopore Technology (ONT) sequencing of four kveik isolates yielded total genome coverage ranging from 300-fold to 1100-fold per sample. We generated both haplotype-collapsed and haplotype-resolved assemblies for each genome. Assembly statistics and BUSCO scores demonstrate that the resulting genomes are of high quality and comparable to the *S. cerevisiae* RefSeq reference genome (Supplementary Data 10). Haplotype-resolved assemblies indicate that the kveik nuclear genome is tetraploid, with an average of 4.4 phased haplotype clusters per chromosome (sample 6). Signatures of strain-specific aneuploidy were also observed, suggesting ongoing chromosomal variability in individual strains (Supplementary Figures 5-8). We found reduced numbers of haplotype clusters covering chrI in three of four samples. Furthermore, sample 7 showed overall less clusters per chromosome and sample 3 showed an increased number of clusters.

### Structural variations predominantly affect repetitive genetic elements

We characterized the structural variations (SVs) in four kveik isolates using an ensemble approach, allowing us to obtain a reliable set of 111 large (≥50bp) regions with insertions and 231 with deletions in at least three of the four isolates. We identified a subset of 217 genes affected in all four isolates. The most prominent class of genomic features impacted by SVs includes transposable elements, long terminal repeats and other repeat sequences (Supplementary Table 1, Supplementary Data 3). Notably, we did not detect large chromosomal rearrangements or fusions compared to the *S. cerevisiae* reference genome.

### Kveik forms an early-diverging paraphyletic group in relation to other European lineages

To uncover the origin of kveik, we performed population genomic analyses using a variety of tools and panels. First, phylogenetic analyses grouped all our kveik taxa into two major geographic clusters (Figure 1): a northern group extending mainly along the Nordfjord region towards Ålesund, and a southern group extending through the Voss/Vossestrand region east of Bergen. Two smaller groups comprised cultures from the Lærdal and Odda regions. Interestingly, cultures from the southernmost region (red) were genetically most alike the northernmost group (green), suggesting possible historical connectivity between these geographically separated areas.

Next, we generated maximum-likelihood phylogenies of the nuclear and mitochondrial DNA (mtDNA) for 302 *S. cerevisiae* (Figure 2). Among the 64 isolates, 62 consistently formed a sister group to Asian fermentation strains within the *S. cerevisiae* nuclear phylogeny, showing strong geographic clustering and minimal within-batch variation (Figure 1, Figure 2A). In order to increase coverage and to unravel the position of kveik yeast in relation to most other wild and domesticated groups, we also generated nuclear maximum likelihood phylogeny comprising 1,269 and 1,307 sequenced genomes from multiple recent studies including two studies on farmhouse yeasts (Bircham *et al*., 2026; Preiss *et al*., 2024; Tellini *et al*., 2024)(Supplementary Figure 2A&B, Supplementary Data 9). Contrary to previous findings (Preiss *et al*., 2024; Preiss *et al*., 2018), our results support an ancient divergence of kveik from other lineages with little evidence for recent gene flow from other populations. When trees were rooted using wild isolates from primordial Chinese forests as outgroup, Asian fermentation and sake appeared closest to the root of the *S. cerevisiae* tree in agreement with previous studies (Duan *et al*., 2018; Peter *et al*., 2018). For the nuclear genome analysis, kveik presents as a paraphyletic group close to the root of domesticated clades, indicating that they represent an early offshoot in the domestication history of *S. cerevisiae*. Mitochondrial phylogenies (Figure 2B) largely mirrored the nuclear genome results, recovering similar clustering patterns for major fermentation groups (e.g. kveik, sake, beer, wine and Asian Fermentation). However, African beer and Dairy Products occupied distinct and divergent mitochondrial lineages, highlighting potential differences in domestication pathways or host environments. Also here, kveik formed an early diverging paraphyletic group in the mitochondrial phylogenetic tree, further supporting the hypothesis that kveik strains represent multiple lineages predating the emergence of industrial beer strains.

### Pronounced population structure in kveik may affect analyses based on allele frequencies

ADMIXTURE analysis revealed pronounced internal population structure within kveik. Five or six distinct allelic clusters can be observed, depending on whether HC2 is further subdivided into two groups corresponding to clusters e and f, (Figure 1B, Figure 3A&B). These clusters closely mirror the phylogenetic groupings and suggest limited but notable outcrossing between regional kveik populations into the HC4 and HC10 clusters. The observed structure implies the presence of long-standing, predominantly regionally isolated ancestral lineages, each comparable in distinctiveness to other major *S. cerevisiae* groups such as beer, sake, and wine.

**Figure 3.**
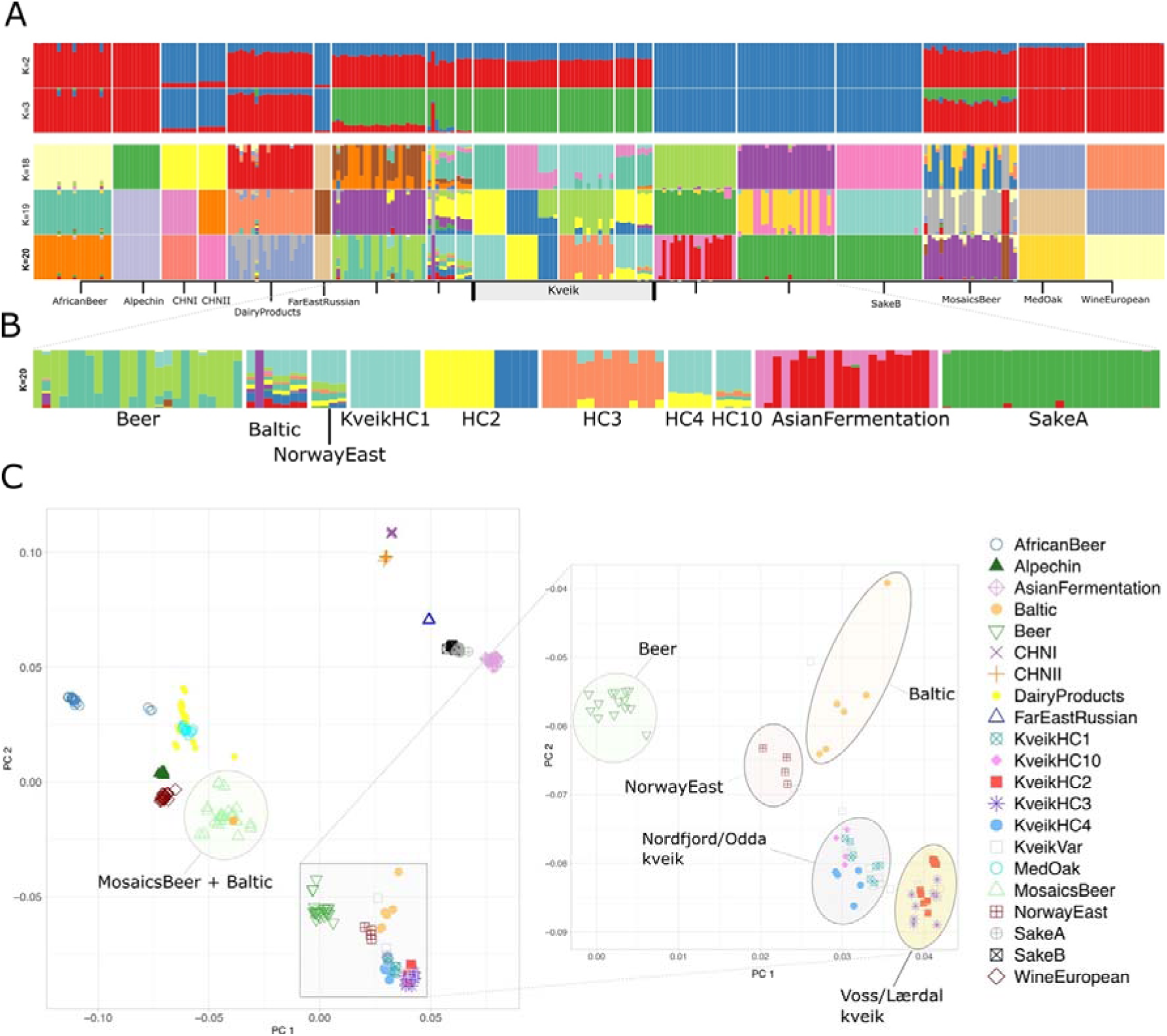
Population structure of kveik, Baltic, eastern Norwegian and selected isolates from the panel compiled by Tellini et al. (2024). A) ADMIXTURE bar plot showing allele contributions for different numbers of clusters (K). Kveik cluster designations are the same as in Figure 1A. Complete ADMIXTURE output is given in Supplementary Figure 2. B) Enlarged ADMIXTURE plot for optimal parameter K=20 yielding minimal cross-validation error using selected populations. C) Projection of variant data onto the first two principal components after principal component analysis. Prominent clusters have been annotated with ellipsoid shapes.

Principal component analysis (PCA) further resolved the geographic structure, placing all kveik isolates into two major subgroups: one predominantly arising from cultures from the Nordfjord region and the other from the Voss region (Figure 3C). These kveik subgroups are adjacent to, but remain genetically distinct from, both the Baltic and eastern Norwegian groups. Notably, one Baltic isolate (Pundurs 1) consistently grouped within the mosaic Beer cluster. The five identified homogeneous clusters (HC) —kveikHC1, −2, −3, −4, and −10 — form well-defined branches in our phylogenetic trees and correspond to distinct ancestral groups in the ADMIXTURE analyses. Isolates exhibiting within-culture variation were grouped into a separate cluster designated KveikVar (Supplementary Figure 2), the ADMIXTURE analysis showing all isolates including KveikVar is depicted in Supplementary Figure 3. Except for kveikHC10, which shows minor allele sharing with the Beer group (Figure 3B) and a single mosaic-like isolate assigned to KveikVar, no significant admixture with external lineages was detected among the kveik clusters. In contrast, the Baltic and, to a lesser degree, eastern Norwegian isolates first described by Preiss *et al*. (2024) exhibit a pronounced mosaic structure, combining gene flow from different western Norwegian lineages with alleles from both Beer and Asian Fermentation groups.

This observation is in accordance with genome-wide genetic distances based on pairwise F_ST_ estimates (Supplementary Figure 4), which reveal high genetic differentiation between Sake and kveik/beer groups, except for the NorwayEast group. These results support the interpretation that kveik has evolved in relative isolation. Incorporating Baltic and eastern Norwegian strains into TreeMix and Bayesian analyses yield a population structure that suggests an early divergence from kveik and supports the main clades assigned (Figure 1, Figure 4).

**Figure 4.**
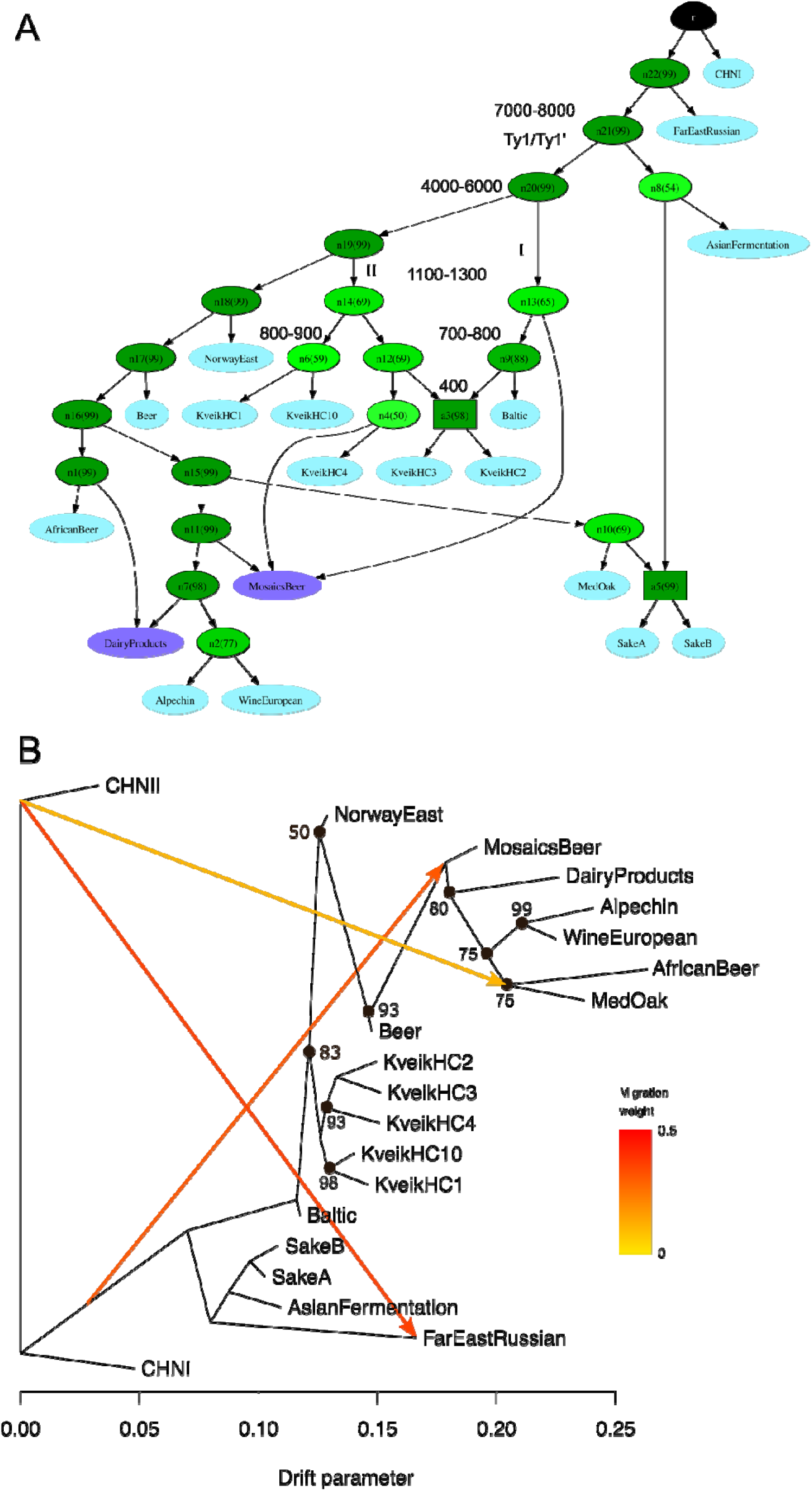
Inferred population history of kveik and other yeast strains by two different methods. A) Admixture graph structure as predicted by AdmixtureBayes including nodes with at least 50% support. Ancestral nodes are plotted in green, selected nodes leading to kveik are annotated with divergence time estimates are rounded to 100 or 1000 years. Node support as percentage of iterations supporting the node is given in parentheses. Leaf nodes in dark blue indicate admixed nodes. Ty1/Ty1’: hypothesized earliest presence of both Ty1 sub-families in an ancestral population; I & II: two hypothesized migration events leading to the introduction of kveik. B) TreeMix tree graph with 3 migration edges with 100 bootstrap iterations. Bootstrap support is given for all nodes having <100% support. Both graphs are rooted using CHNII as outgroup.

Previous TreeMix analyses had pooled Norwegian and Baltic strains into a hypothetical “European Farmhouse” population. There is, however, evidence for a pronounced structure (and possibly admixture in the Baltic group) in these analyses (compare Figure 4a, cluster 11B “European Farmhouse” in Preiss et al. 2024) showing a strong migration signal between kveik and the root of Asian fermentation cluster (compare Figure 5a in Preiss et al. 2024). While we were initially able to produce a similar plot (Supplementary Figure 11), this signal disappears when the Norwegian and Baltic strains are assigned to individual clusters (Figure 4B). Importantly, while the revised TreeMix and Bayesian models show no gene flow between kveik and Asian strains, both detect a strong signal of gene flow from diverse sources into the Mosaic beers group in accordance with prior expectations (Figure 4A). This highlights the complexity of interpreting admixture signals and underscores the importance of accounting for internal structure when modeling yeast population histories.

**Figure 5.**
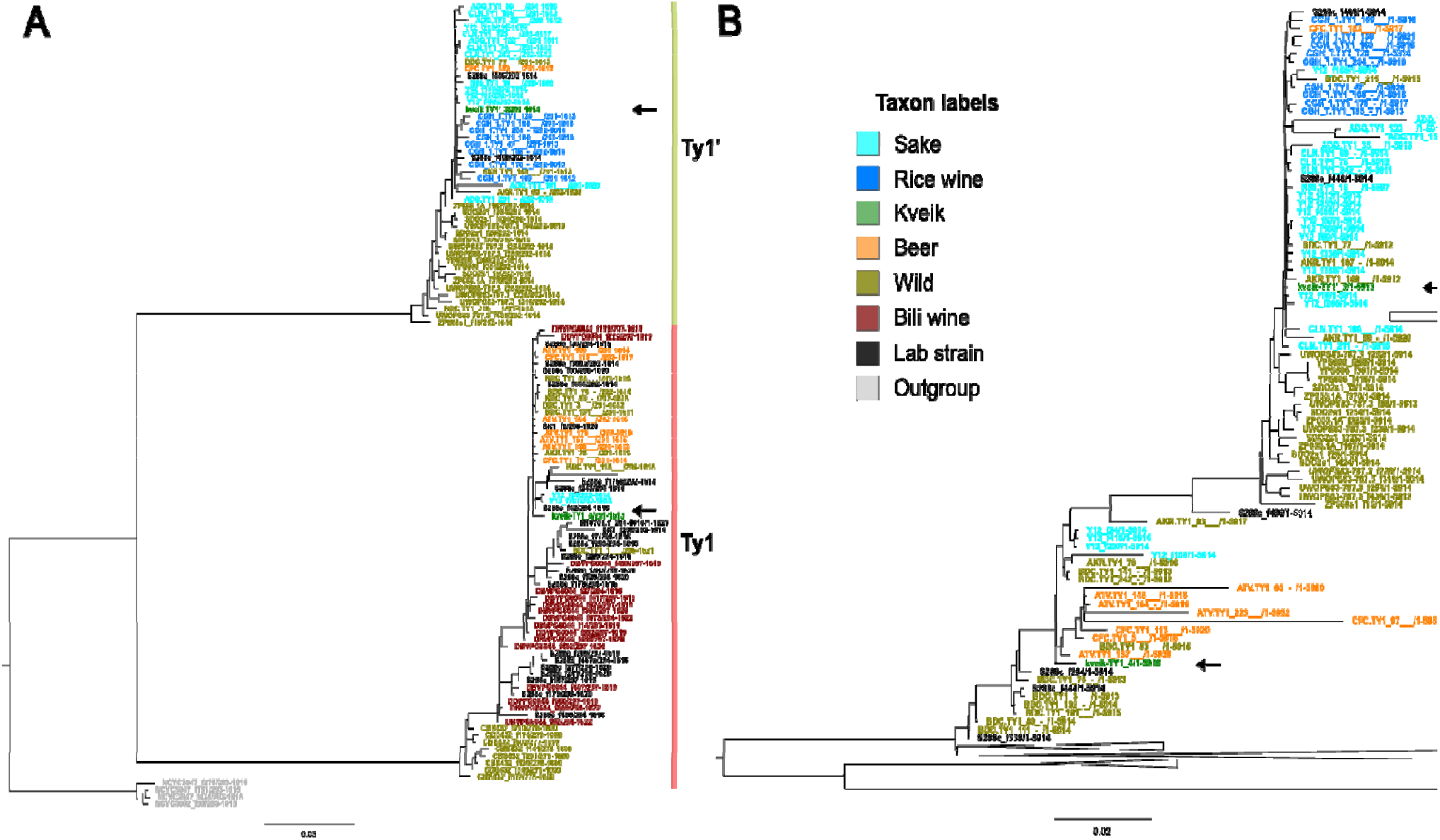
Maximum-likelihood phylogenies of Ty1-like TEs in kveik assemblies. Comparison to the panel from Czaja et al. (2020) includes homologous kveik sequences (in green) from our long-read assemblies and additional Ty1 elements extracted from recent assemblies (O’Donnell et al., 2023). A) Phylogeny of the gag-genes based on 307 parsimony-informative sites. The S. cerevisiae Ty1 family of retrotransposon can be divided into two groups based on their gag elements: canonical Ty1 (red) and Ty1’ (green). Kveik genomes contain a single full-length Ty1 element of each class. B) Phylogeny of full-length Ty1-elements including kveik sequences based on 1701 parsimony-informative sites. Taxon label color codes are the same in A) and B). Both trees were rooted on the Saccharomyces jurei (grey) clade. Positions of kveik taxa are indicated by arrows. All homologous sequences extracted from our four assemblies had 100% sequence identity and are therefore represented by a single entry.

### Simulation experiments show TreeMix analysis can be affected by pooling

Admixture signals between sake and beer strains have been previously reported based on coalescent theory, prompting us to investigate whether the migration signals detected in our data could be confounded by the specific demographic histories of the strains involved. Modern sake strains are thought to have passed through a strong and recent population bottleneck (Ohya & Kashima, 2019). To explore this possibility, we ran extensive simulations using a variety of demographic models, incorporating divergence times and evolutionary rates based on existing literature. These models included scenarios with and without admixture as well as populations with bottlenecks of varying strengths, and more complex internal population structure similar to kveik (Supplementary Figure 9). The results from TreeMix indicated migration signals for both models with and without admixture. False-positive migration signals were observed in demographic models that included a single strong bottleneck for the simulated sake population and multiple weaker bottlenecks for the simulated kveik population. A similar effect was also observed in a more complex model when a simulated ‘Kveik’ group consisting of multiple isolated populations was analyzed as a single pooled population, but not when the ‘Kveik’ group was subdivided (Supplementary Figure 10, Supplementary Note Admixture simulation). This confirmed our hypothesis that migration signals can readily be evoked by bottlenecks or more generally may arise as pooling artifacts through unaccounted population structure in collapsed clades.

### Transposable element inventory shows limited expansion

The only families of transposable elements (TE) believed to be active in extant *S. cerevisiae* lineages belong to the families of Ty1 and Ty2 Long Terminal Repeat (LTR) retrotransposons. RepeatMasker searches identified orthologous full-length Ty-sequences in low abundances. Among all assemblies investigated, kveik exhibit some of the lowest copy numbers of complete Ty elements (Supplementary Data 4). Specifically, we identified five hits to the internal protein-coding domain of the Ty1 family, of which three were truncated and only two appeared to be complete. Importantly, all Ty1-like sequences were identical in both sequence and locus across all four kveik genomes, allowing us to analyze them jointly. Phylogenetic analysis of the gag protein sequences places them in the stem group of Ty1’ elements, while one is placed jointly with canonical Ty1-gag genes from (Figure 5). Based on these findings, we refer to the full-length elements as kTy1 (canonical) and kTy1’ (derived) and propose three likely incomplete ancestral transposition events for kTy1’, and none for kTy1. Notably, kTy1’ forms a clade with the Ty1’ sequences from Sake strain Y12, suggesting an intriguing phylogenetic connection.

### Divergence time estimation depends on specifics of the traditional brewing process

We estimated divergence times between kveik and other *S. cerevisiae* lineages based on pairwise differences at 4-fold degenerate sites, similar to a previous approach using mutation rates and generation times found in the literature (Gallone *et al*., 2018; Peter *et al*., 2018). The largest median internal divergence time between different kveik strains depending on mutation rates was estimated to be 1046 to 1152 years ago (kveikHC1 and −HC3), 737 to 1332 years ago between kveik and the Baltic strains, and 1528 to 2310 years ago between kveik and the strains from eastern Norway. Groupwise estimates show high variability between strains. An overview of the estimated divergence times and confidence intervals for the groups of strains are given in Table 2.

**Table 2.** Pairwise divergence time estimates between kveik and other selected groups. Time estimates are given in years rounded to the nearest integer, based on 1,128,220 4-fold degenerate sites across 254 samples assuming G_1_ = 2920 (Peter et al. 2018) and G_2_ = 30 generations/year and mutation rates µ_1_ =1.84 ・10^−10^, µ_2_ = 1.67 ・10^−10^ (Peter et al., 2018), µ_3_ = 1.61 ・10^−8^(Gallone et al., 2016). Extreme values within the kveik group are set in bold. *: admixture may affect estimates in the Baltic group.

| Group 1 | Group 2<br>(KveikHC) | $\mu_1$, G<sub>1</sub> | $\mu_2$, G<sub>1</sub> | $\mu_3$, G<sub>2</sub> | Median gen. $\mu_1$ | Median gen. $\mu_2$ | Median gen. $\mu_3$ |
| --- | --- | --- | --- | --- | --- | --- | --- |
| KveikHC1 | **2** | 1031 | 1136 | 1147 | 3.01182e+06 | 3.318413e+06 | 3.44208e+04 |
| KveikHC1 | **3** | 1046 | 1152 | 1164 | 3.055205e+06 | 3.366214e+06 | 3.491663e+04 |
| KveikHC1 | **4** | 418 | 461 | 465 | 1.221373e+06 | 1.345705e+06 | 1.395855e+04 |
| KveikHC1 | **10** | 809 | 892 | 900 | 2.363487e+06 | 2.604082e+06 | 2.701129e+04 |
| KveikHC2 | **3** | 388 | 427 | 431 | 1.132183e+06 | 1.247435e+06 | 1.293923e+04 |
| KveikHC2 | **4** | 251 | 276 | 279 | 7.322696e+05 | 8.06812e+05 | 8.368795e+03 |
| KveikHC2 | **10** | 457 | 504 | 508 | 1.334564e+06 | 1.470417e+06 | 1.525215e+04 |
| KveikHC3 | **4** | 398 | 438 | 442 | 1.161094e+06 | 1.279289e+06 | 1.326964e+04 |
| KveikHC3 | **10** | 542 | 598 | 604 | 1.585144e+06 | 1.746506e+06 | 1.811593e+04 |
| KveikHC4 | **10** | 323 | 356 | 360 | 9.442669e+05 | 1.04039e+06 | 1.079162e+04 |
| Baltic* | **1** | 1209 | 1332 | 1345 | 3.530076e+06 | 3.889425e+06 | 4.034373e+04 |
| Baltic* | **2** | 737 | 812 | 820 | 2.151416e+06 | 2.370423e+06 | 2.458761e+04 |
| N. East | **1** | 2097 | 2310 | 2332 | 6.122351e+06 | 6.745584e+06 | 6.996973e+04 |
| N. East | **2** | 1528 | 1683 | 1699 | 4.460695e+06 | 4.914778e+06 | 5.097937e+04 |
| Beer | **1** | 3069 | 3382 | 3414 | 8.962483e+06 | 9.874831e+06 | 1.024284e+05 |
| Beer | **2** | 2440 | 2689 | 2715 | 7.125944e+06 | 7.851339e+06 | 8.143936e+04 |
| *Sake A* | **1** | 8605 | 9481 | 9572 | 2.512636e+07 | 2.768414e+07 | 2.871584e+05 |
| *Sake B* | **1** | 9026 | 9945 | 10041 | 2.635766e+07 | 2.904078e+07 | 3.012305e+05 |
| *Asian Ferment.* | **1** | 7670 | 8451 | 8532 | 2.239604e+07 | 2.467588e+07 | 2.559547e+05 |
| *CHN I* | **1** | 7396 | 8149 | 8227 | 2.159691e+07 | 2.37954e+07 | 2.468219e+05 |
| *CHN II* | **1** | 7118 | 7842 | 7917 | 2.078343e+07 | 1.810049e+07 | 2.375249e+05 |

## Discussion

We have sampled and sequenced the largest collection of kveik original brewing cultures from traditional farmhouse brewing sites to date, aiming to gain insights into the demographic history and reconstruct the present haplotypes of domesticated *S. cerevisiae* populations. Variation within domesticated yeast cultures has been largely overlooked in previous research. Our study is one of the first studies that included multiple isolates from brewing cultures, allowing us to assess genetic strain variation within them. While we observe variation, predominantly in those yeast brewing cultures we assigned to the KveikVar group, a majority of isolates cluster closely together by original culture and geographical region. Compared to commercial monocultures, genetic diversity was unsurprisingly higher in kveik, even though some of the isolates had low internal diversity comparable to commercial strains.

Bircham *et al*. (2026) recently profiled genetic diversity across 44 farmhouse brewing cultures from regions that partially overlap our sampling, using fingerprinting of individual isolates and Illumina sequencing for a subset. They report that many farmhouse cultures comprise mixtures of strains, with highly variable within-culture diversity and some geographic structuring – patterns that are broadly consistent with our own results and supportive of a complex domestication and dispersal history for kveik. Their genotyping further indicates that Baltic and eastern Norwegian isolates are genetically distinct from the kveik lineage, with admixture/introgression signals concentrated in the Baltic group, reinforcing our conclusion that current evidence does not support a single, pan-European “farmhouse” population that would unite these groups with kveik.

Taken together, these studies support and further refine earlier hypotheses that individual traditional brewing cultures might represent a complex population of different strains (Preiss *et al*., 2018). Future research should investigate how genetic diversity shapes, and is shaped by, the unique ecology of traditional brewing strains.

Norwegian and Baltic isolates showed the highest levels of heterozygosity among all, which may indicate healthy and robust populations maintained by genetic diversity. On the other hand, heterozygosity may affect genome assembly continuity and can slightly inflate branch lengths in assembly-based phylogenies, although it is not expected to alter phylogenetic topology (Lischer *et al*., 2014).

Previous research has indicated discordant evolutionary histories for yeast nuclear and mitochondrial genomes of *S. cerevisiae* (De Chiara *et al*., 2020). In this study, we find congruence between both nuclear and mitochondrial genomes of 302 *S. cerevisiae* strains, albeit with some important exceptions in the African beer and dairy products groups. However, there are signs of accelerated mitochondrial evolution within some of the kveik subgroups.

Based on phylogenetics and divergence time estimates, we hypothesize that the common ancestor of beer, kveik, and other domesticated lineages emerged from an ancestral population between 4000 to 8000 years ago, undergoing vastly different domestication histories thereafter. Our estimate also coincides with the earliest documented emergence of barley in neolithic China and first evidence of mixed beer fermentation by the Yangshao people approximately 5000 years ago (Wang *et al*., 2016).

Phylogenetic analyses of kveik genomes and mitochondrial genomes all indicate an early divergence of the kveik group, possibly following human migration and the spread of agriculture (Peña *et al*., 2025). Our results suggest that kveik is paraphyletic and a sister group to Asian fermentation within the domesticated lineages, whereas previous reports have depicted kveik as a single compact group implying monophyly, likely due to the limited number of samples included.

The pronounced population structure unveiled by our comprehensive approach likely results from geographical isolation, combined with the ecological niche created by long-standing brewing traditions shaping a group of strains autochthonous to western Norway. Furthermore, geographic distribution and Bayesian analyses led us to hypothesize a complex migration history of kveik with multiple distinct founders into vastly separated landscapes, largely coinciding with the historical regions of Hordaland and Firdafylke under the older Gulating jurisdiction.

We speculate that western Norwegian lineages descended from ancestral populations at a minimum 800 – 1300 years ago. However, in the absence of fossil records for calibration, such estimates must be treated with extreme caution. Despite this, most biases introduced by e.g. variant filtration and generation times, will possibly lead to systematically underestimate divergence times. The number of generations per year is of relevance in this context, given the unique tradition of farmhouse brewing. Codified in the older Gulating law (Simensen, 2021) and also mentioned in older sources, medieval Norwegian society mandated brewing at least twice per year predating Christianization in the 11^th^ century. If traditional brewing practices have been largely preserved over an extended period, the practice of using kveik for only a few, short brewing sessions—after which it is dried—may result in lower numbers of generations per year by an order of magnitude, and thus older time estimates.

The availability of T2T assemblies of S. cerevisiae has enabled precise inference of their transposon copy numbers (Czaja *et al*., 2020; O’Donnell *et al*., 2023). It has been shown that TE repertoires are lineage-specific across *S. cerevisiae* populations (Bleykasten-Grosshans *et al*., 2013) and this is also evident for kveik. Here, we used the SCRaP panel of T2T assemblies (O’Donnell *et al*., 2023) to enable direct comparison of full-length, truncated, and solo-LTR elements, allowing explicit consideration of alternative explanations for reduced TE burden. Kveik assemblies showed unusually low total Ty copy numbers relative to most domesticated strains. This pattern may reflect limited TE expansion, ecological constraints associated with traditional farmhouse brewing, or historical demographic effects. Secondary TE loss is another possible explanation; however, we did not observe an enrichment of solo-LTR elements expected under extensive deletion. Recent research has focused on the mechanisms behind the Ty1 Copy Number Control (CNC) system (Cottee *et al*., 2021; Garfinkel *et al*., 2003; Garfinkel *et al*., 2016), though CNC in *S. cerevisiae* remains incompletely understood (Ahn *et al*., 2017; Czaja *et al*., 2020). The low copy number of Ty elements in kveik could be of particular interest in future studies exploring the evolutionary mechanisms regulating retro-transposition.

Previous studies have suggested secondary gene flow from Asian fermentation or sake strains into European lineages (Fay *et al*., 2019; Preiss *et al*., 2024; Saada *et al*., 2022). However, our analyses did not find sufficient support for gene-flow into western Norwegian lineages. Several factors may have contributed to previous discoveries of admixture signals. A key methodological difference between this study and earlier works is how kveik or farmhouse yeast was treated either as a single group or multiple coalescent. By improving strain representation, we detected sufficient internal population structure, notably in accordance with results presented by Preiss *et al*. (2024), to justify subdividing it into clusters that may have formed because of regional isolation and genetic drift. It has been established that recent population bottlenecks can cause spurious admixture signals in population structure analyses (Lawson *et al*., 2018). We demonstrate that a similar “phantom-migration” effect can be observed in TreeMix on data simulated under a coalescent model with no admixture, but with population bottlenecks of varying strengths. Other contributing factors may have been copy number expansions of identical Ty element alleles between kveik, sake, and the reference genome, and the inclusion of sake-like or mosaic beer strains (Preiss *et al*., 2024).

Considering the possible Far Eastern or Eastern European origin of kveik, its apparent endemism to western Norway is a paradox. In accordance with previous results, Baltic and, to a lesser degree, Eastern Norwegian strains show evidence for mosaicism with gene flow from western Norwegian lineages and other sources (Preiss *et al*., 2024); consequently this finding could indicate more recent hybridization or admixture events. If these are genuine, admixture signals may themselves help to resolve more recent migration histories of strains from Norway into other regions.

We have focused our aneuploidy, copy-number and broader SV analyses on the four kveik genomes with high-confidence long-read sequences and T2T assemblies due to their ability to resolve repetitive and particularly sub-telomeric regions (O’Donnell *et al*., 2023) whereas phylogenetic reconstruction relied on short read assemblies. The potential impact of copy-number alterations and gene loss or gains on the phylogenetic signal have not been part of this investigation. While the majority of SVs detected by our pipeline were associated with TEs, complex unresolved aneuploidies may exist in kveik. Previous studies did, however, not clearly indicate that patterns of aneuploidy are population specific (Duan *et al*., 2018; O’Donnell *et al*., 2023). Testing population-level specificity of aneuploidy and SV would require broader long-read sampling and is beyond the scope of the present study.

The erosion of cultural practices and replacement of traditional by industrial strains can lead to loss of genetic diversity in domesticated *S. cerevisiae*. While we found little evidence for ongoing admixture or hybridization with industrial strains, the replacement of traditional strains or the creation of hybrid strains for research and new products remains a possibility. The growing popularity of traditional brewing strains and their widespread distribution by commercial retailers could help preserve traditional strains but will likely also distort the local population structure that evolved in isolation over centuries. The data generated in our study contribute to preserving these ancient haplotypes as representatives of an early-diverging domesticated lineage of *S. cerevisiae*.

## Conclusions

Our study establishes traditional Norwegian kveik yeast as an early-diverging, paraphyletic lineage within domesticated *S. cerevisiae*. The survival of yeast strains may be closely linked to the intangible cultural heritage of farmhouse brewing, which has survived in isolated locations such as western Norway. By disentangling true population history from admixture artifacts, we highlight the importance of accounting for internal population structure and selection. These findings reshape our understanding of yeast domestication and demonstrate how traditional, community-maintained strains can serve as living archives of evolutionary processes. Our work also underscores the value of preserving such lineages as part of both biological and cultural heritage.

## Materials and Methods

A graphical display of materials and methods is given in Figure 6. Graphical overview of the samples and analysis workflow used in this study. The term “original brewing culture” (or “culture” in short) is used for the original yeast samples from traditional brewing (liquid or dry), the term “isolate” is used for the 64 sequenced genomes isolated from single colonies, and the term “strain” is used for a unique genomic lineage represented by one or more isolates.

**Figure 6.**
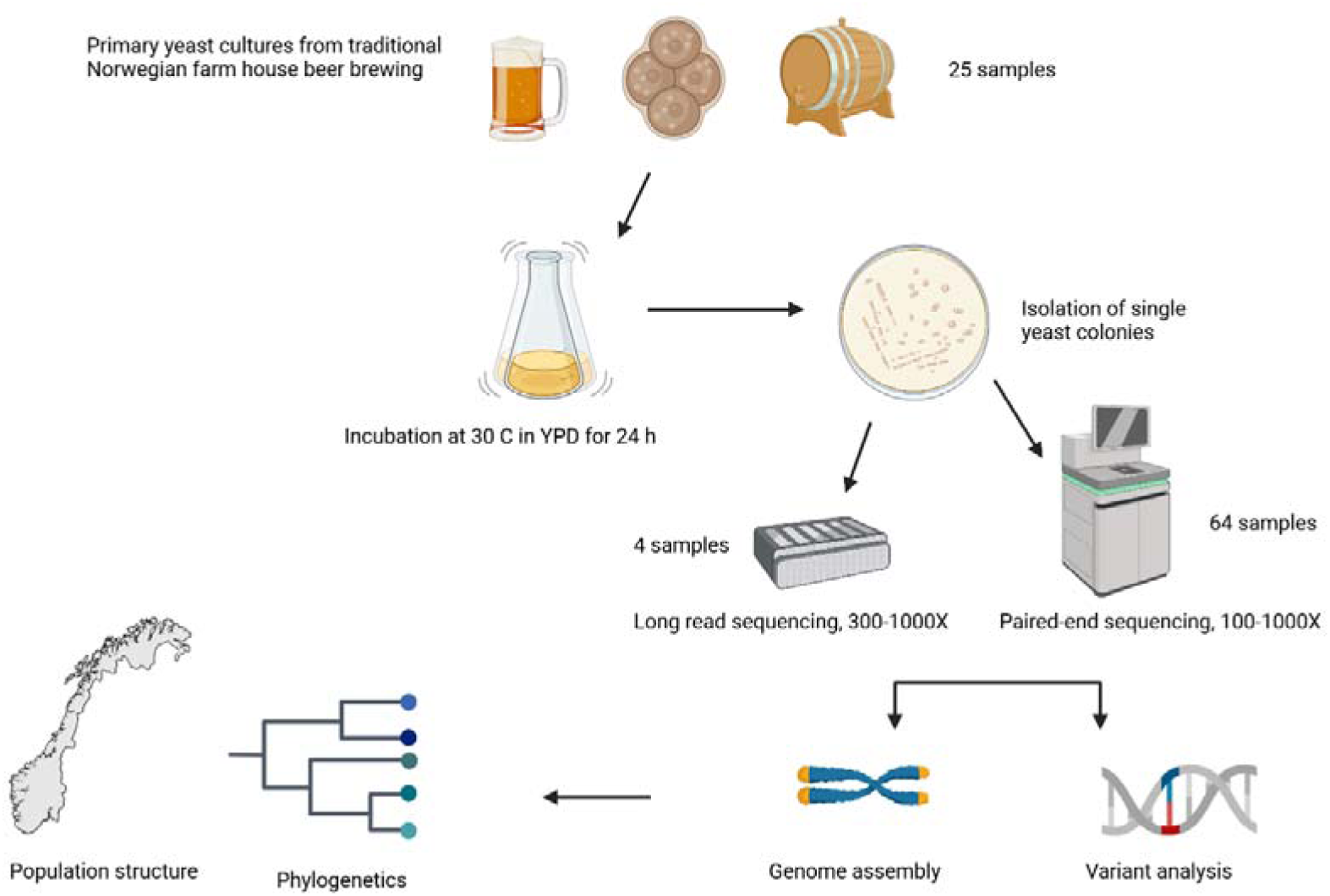
Graphical overview of the samples and analysis workflow used in this study.

### Sampling

Traditional farmhouse brewing has long traditions in western Norway and is still actively practiced. Through knowledge of these traditions and networks in the region we were able to locate brewers and document brewing traditions from previous generations. Key criteria for sampling were: 1. Kveik had been in the family for several generations. 2. The intangible cultural heritage connected to brewing with kveik was still practiced (see Garshol (2023)). 3. The yeast had recently been used for brewing or could be traced back to brewing traditions at the location, and the owners were willing to be interviewed regarding the processes of brewing and the use of kveik yeast. Samples (n = 25) were collected directly from the owners with a written contract granting use in research (Table 1and Figure 6). Sample 1 was re-sampled after two years from the same owner (sample 1A and 1B), and sample 3 and 27 represent a new (2020) and an old sample (barrel > 50 years old) from the Selland farm, respectively. Multiple single colonies (2-4) were isolated from 17 different kveik samples subjected to genomic sequencing (Supplementary Data 2).

### Yeast cultivation and single colony isolation

Dried kveik samples were rehydrated in sterile water. The rehydrated samples of kveik and the liquid yeast slurries were enriched by inoculating 100 μl of the slurry into 100 ml YPD medium (1% yeast extract; 2% peptone; 2% dextrose; chloramphenicol 0.3 g/l). The samples were incubated at 30□ for 48 h while shaking, then streak plated onto Wallerstein Nutrient Agar (with chloramphenicol 0.3g/l) (WLN; Thermo Fisher Scientific, CM0309), a differential medium for yeasts that can distinguish multiple strains within one sample based on uptake of the bromocresol green dye. Yeast colonies were then sub-cultured twice onto WLN to ensure purity. The single colonies were enriched for DNA extraction as described below. The remaining overnight cultures were long-term stored in aliquots at minus 80□ using a glycerol-based standard storage medium (peptone 1% w v-1, yeast extract 0.5% w v-1, glucose 1% w v-1, glycerol 25% v v-1).

### DNA extraction

DNA was released from yeast cells in an aqueous phase using a modification of previous protocols (Ausubel, 2002; Preiss *et al*., 2018) and then directly followed by DNA extraction applying the Qiagen DNeasy Plant Mini kit. Briefly, yeast cells were collected from 1.5 ml of the overnight yeast culture by pelleting cells with centrifugation at 13,000 rpm for 5 min, removing the supernatant, washing the cell pellet in 1 mL of 0.8% NaCl and adding 0.3g of 0.5-mm-diameter glass beads. The cell-bead mixture was centrifuged at 11,600 g for 10 min at 4 °C, and the supernatant was discharged followed by resuspension in 300 μl of breaking buffer (2% Triton X-100, 1 % SDS, 100mM NaCl, 10 mM Tris-HCl) and 300 μl of phenol-chloroform-isoamyl alcohol (25:24:1). The suspension was vortexed at high speed 3 x 45 sec, centrifuged at 11600 x g for 10 min at 4 °C, and the final aqueous phase was then used for DNA extraction with the DNeasy Plant Mini Kit (Qiagen, Tokyo) according to the manufacturer’s instructions. High-molecular weight (HMW) DNA was extracted from 4 ml of overnight yeast culture using the Qiagen Genomic-tip 20/G according to the manufacturer’s instructions. Prior to loading on the genomic-tip columns, lysis of yeast cells was performed with 100μl Lyticase (1000 units/ml, Sigma Aldrich) as described in the Qiagen Genomic DNA Handbook (release date: 06/2015). DNA quantity and length were tested on a Nanodrop spectrophotometer and a 0.6% Agarose gel (> 50 kb).

### Short read genome sequencing

DNA samples were quantified using the Qubit fluorometer and 100ng of each sample was used for library construction with Illumina TruSeq DNA nano prep. Quantification of libraries was carried out by using KAPA Library Quantification kit (qPCR) for Illumina, and average library input fragment size was estimated using an Agilent TapeStation. Sequencing was performed on the Illumina NovaSeq 6000 platform using standard workflow and sequencing parameters, employing paired-end sequencing (read length 2 x 150bp) with an average depth of over 400-fold for each sample.

### Long read sequencing

DNA was stored at 4°C for extended periods of time to avoid freeze-thaw cycles. To quantify HMW-DNA, an aliquot of the HMW DNA was homogenized using a glass bead and measured in triplicates on a Nanodrop One (Thermo Fisher Scientific) and once on a Qubit (Thermo Fisher Scientific), following the methods described by Koetsier & Cantor (2021). The integrity of the HMW DNA was assessed by running the DNA on a 0.6% agarose gel at 3V/cm in 0.5xTBE buffer and alongside a 1kb extent DNA ladder (NEB) with an upper band of 48.5kb. DNA was processed for sequencing if the main band was above 48.5kb and no smear was visible below 20kb.

Nanopore Library preparation was performed using a LSK114 ligation-based sequencing kit (Oxford Nanopore) according to manufacturer’s instructions with minor modifications. Briefly, HMW DNA was used at a concentration of 20-50ng/µl, with a total of 1-2.5µg per sample. When the concentration of DNA was lower than 20ng/µl, the initial end preparation reaction volume was scaled up (i.e. 2x volume for 10ng/µl) to reach a total DNA amount of at least 1µg per sample. To avoid shearing of DNA by pipetting, wide-bore tips were used, and DNA was added directly to premixed reaction mixes, where possible. To maximize library recovery, the final elution was performed with 25µl, and incubation time was extended to 30 minutes at 37°C. Final libraries were quantified on a Qubit 3 fluorometer (Thermo Fischer Scientific) and stored at 4°C until sequencing.

For sequencing, a R10.4.1 PromethION flow cell was primed according to manufacturer’s instructions and sequencing was performed on a P2Solo device (Oxford Nanopore) with live base calling enabled (MinKnow version 23.11.4). The flow cell was washed after each sequencing run using the flow cell wash kit (Oxford nanopore) according to manufacturer’s instructions and stored at 4°C. All sequencing runs were completed within two weeks. Base calling was performed using dorado (version 0.5.3) with the inherent dna_r10.4.1_e8.2_400bps_sup@v4.2.0 model, resulting in a read N50 between 23 and 33kb and a mean Q-score of 18.6-20.3. For sample 6 that was sequenced to greater depth candidate duplex reads were identified using the dorado duplex tool suite (version 0.5.3) and duplex base calling was performed using the dna_r10.4.1_e8.2_5khz_stereo@v1.1 model, resulting in a mean Q-score of 28.0 and a read N50 value of 23,229bp.

### Genome assembly

Illumina sequences were quality checked and trimmed with fastp (Chen *et al*., 2018). Genome assemblies for strains sequenced by Illumina only were generated with SPAdes (v4.0.0) (Bankevich *et al*., 2012). A haplotype-collapsed high-quality genome assembly was derived from sample 6. Long reads were first filtered to minimum length 10kb and quality > Q20 with Seqkit (Shen *et al*., 2024) and processed with Porechop. Simplex reads were initially assembled with Flye (v2.9.3-b1797) (Kolmogorov *et al*., 2019) with 4 rounds of polishing, followed by 4 rounds of polishing with Medaka (1.11.3) (Oxford Nanopore Technologies) using duplex reads, and one round of polishing with Pilon (v1.11.3) (Walker *et al*., 2014) using the Illumina reads from the same sample. The resulting contigs were scaffolded with RagTag (Alonge *et al*., 2022). The *S. cerevisiae* S288C RefSeq genome assembly R64-2-1 (GCF_000146045.2) (S288C_R64) was used as reference (Engel *et al*., 2022). Haplotype-resolved assemblies were generated by clustering reads with nPhase (Abou Saada *et al*., 2021) consecutively running on each cluster separately. No further polishing was applied to these assemblies to avoid masking heterozygous variants. The quality of the resulting assemblies was assessed with Quast (Mikheenko *et al*., 2018), BUSCO (using OrthoDB saccharomycetaceae_odb12) (Manni *et al*., 2021), and D-GENIES (Cabanettes & Klopp, 2018).

### Genome annotation

The assembled genomes were annotated for various genomic features, including protein-coding genes, tRNAs, transposable elements (TE), and telomere-associated elements by a custom Snakemake-workflow derived from the LRSDAY-framework (Yue & Liti, 2018). Briefly, centromeric regions in the nuclear genome were identified and annotated using the exonerate tool. Y’ elements were detected with Blat, and core X-elements were detected with HMMER. MAKER was used for nuclear gene prediction and annotation, and EvidenceModeler (EVM) for refining multi-exon gene annotations. For mitochondrial gene annotation, mfannot was used for gene prediction, with the genetic code table for yeast mitochondria, and tRNAscan-SE for identifying tRNA genes. Orthologous genes were identified using the proteinortho tool. An alternative workflow for TE detection and annotation was implemented and applied following a protocol and using repeat libraries described previously (Czaja *et al*., 2020), based on RepeatMasker, RMblast, and custom Perl scripts.

### Variant analysis

The workflow for variant detection was designed to process sequencing data using the Genome Analysis Toolkit (GATK, v4.5.0.0) (Mckenna *et al*., 2010). Raw data from Preiss *et al*. (2018), Gallone *et al*. (2016), and Tellini *et al*. (2024) were downloaded from the Sequence Read Archive (SRA). Quality control and trimming of the raw sequencing reads were performed using fastp. Trimmed reads were aligned to the S288C_R64 reference genome using BWA (v0.7.18), and sorted and indexed using SAMtools (v1.20) (Danecek *et al*., 2021). Duplicate reads were marked using GATK’s MarkDuplicatesSpark. Variants were called using GATK’s HaplotypeCaller in GVCF mode, with ploidy parameter set to 2 and 4 consecutively. The GVCF files were indexed and imported into a GenomicsDB, joint genotyping was performed excluding the mitochondrial genome. Variants were filtered using GATK’s VariantFiltration based on the following exclusion criteria: mapping quality (MQ < 50.00), quality-to-depth ratio (QUAL / DP < 2.0), read depth (DP < 5), and genotype quality (GQ < 30) retaining only biallelic SNPs and InDels. Summary statistics of the variant call files were calculated using BCFtools v1.20, using method bcftools stats and a custom AWK script (Danecek *et al*., 2021). Heterozygosity was defined as: number of heterzygous sites/total number of sites. The R-package PopGenome (Pfeifer *et al*., 2014). mtDNA variants were called by aligning reads to the mitochondrial reference sequence with bwa (Li & Durbin, 2010), alignments were filtered for minimum MQ 30, and variants where called with BCFTools and filtered for minimum DP of 3.

### Structural variant prediction

The trimmed Nanopore reads were aligned to the S288C_R64 reference genome using Minimap2 (v2.24) (Li, 2021), and the aligned reads were sorted and indexed using SAMtools (v1.20). Structural variants (SVs) were detected using multiple tools: cuteSV (v1.0.11) (Jiang *et al*., 2020), Sniffles (v2.0.7) (Smolka *et al*., 2024), pbsv (v2.8.0) (PacificBiosciences), NanoVar (v1.4.1) (Tham *et al*., 2020), SVIM (v1.4.2) (Heller & Vingron, 2019), and NanoSV (v1.2.4) (Cretu Stancu *et al*., 2017) with default parameters. The results from these tools were then combined using combiSV (v2.2) (Dierckxsens *et al*., 2021) into a consensus call-set of SVs. To minimize false positive SV calls, each variant required support from at least two out of six tools. A high confidence set of genes affected by SVs was generated by including genes overlapping with an SV in all four strains.

### Phylogenetic analysis

*Whole-genome alignments of draft assemblies were used to extract parsimony-informative sites for maximum-likelihood phylogenetic inference in IQ-Tree (Fig. 2), with ModelFinder model selection and ultrafast bootstrap support.* Reference-based alignments were performed using the NASP pipeline (Sahl *et al*., 2016) with default parameters using NUCmer to align assembled genomes (Delcher et al. 2003). IQ-Tree2 (v2.3.6) (Minh *et al*., 2020; Nguyen *et al*., 2015) was used with 1000 iterations of the ultrafast bootstrap approximation (Minh et al. 2013) using ModelFinder for model selection by BIC (Kalyaanamoorthy *et al*., 2017). Best-fit models chosen by ModelFinder were TVM+F+ASC+R4 (Figure 1B), TVMe+ASC+R8 (Figure 2A), TVM+F+I+R7 (Figure 2B). ML-phylogenies of 1269 and 1306 SPAdes assemblies (Supplementary Figure 2A&B) were generated as above, but with IQ-Tree3 (v3.0.1) using a preset general time-reversible model (GTR+ASC+G). Phylogenetic trees were visualized in iTol (Letunic & Bork, 2024). Nucleotide sequences of Ty genes (Figure 5) were aligned by MUSCLE (Edgar, 2004) and inspected and manually clipped in Jalview (Waterhouse *et al*., 2009) and ML-phylogenies were inferred with IQ-Tree2 with 1000 ultrafast bootstrap iterations, models selected by BIC: TPM3u+F+I+R2 (Figure 5A), TVM+F+I+R4 (Figure 5B). Phylogenetic trees were visualized with FigTree (Rambaut, 2023).

### Assignment to homogeneous clusters (HC)

To define conservative groups for allele frequency–based analyses, we used the rooted nuclear SNP phylogeny of kveik isolates (Fig. 1B), independent of sampling origin, and partitioned the kveik portion into a minimal set of well-supported monophyletic clades such that each clade contained only kveik taxa. We assigned an original brewing culture to a homogeneous cluster (HC) if, and only if, all its isolates fell within the same clade. Cultures whose isolates were distributed across multiple major clades were classified as KveikVar and excluded from allele frequency–based analyses (ADMIXTURE, TreeMix, AdmixtureBayes) in the manuscript but retained in phylogenetic plots as well as Supplementary Figure 3 for completeness, cultures containing deep-branching outliers (cultures 7, 8, and 18) were conservatively assigned to KveikVar as well.

### Population genomics and admixture

Population genomics analyses were conducted using filtered biallelic SNPs. The primary tools and their versions used in the analysis included: VCFtools (v0.1.16), Plink (v1.90b6.21), ADMIXTURE (v1.3.0) (Alexander *et al*., 2009), AdmixtureBayes (Nielsen *et al*., 2023), TreeMix (v1.13) (Pickrell & Pritchard, 2012), and OptM (version 0.1.6) (Fitak, 2021). SNPs with high linkage disequilibrium (LD) were pruned using a custom script (‘ldPruning.sh’) with an r² value greater than 0.2 within a window of 50 kb, step-size10 kb. The output was converted to TreeMix format using customized shell and python scripts. TreeMix was used to infer population trees and migration events, with 100 bootstrap replicates allowing for 0 to 12 migration events. Different numbers of migration events were tested and optimized using OptM. Phylip consense (version 3.697) was used to generate consensus trees. ADMIXTURE analysis estimated individual ancestry proportion of ancestral populations (K = [2,22]) and was visualized using a customized R script. AdmixtureBayes was used to run MCMC simulations with 1 million iterations, and convergence was evaluated using trace plots, Gelman-Rubin diagnostics, and autocorrelation plots. Pairwise F_ST_ values were estimated with AdmixTools 2 (Maier *et al*., 2023) and plotted with the ComplexHeatmap package in R.

### Admixture simulation

Genetic data was simulated with msprime (version 1.3.3) (Baumdicker *et al*., 2022) and variants were analyzed with TreeMix as described above. Details on the demographic models used are provided in Supplementary Information Admixture Simulation.

### Divergence time estimation

4-fold degenerate sites were annotated with degenotate (https://github.com/harvardinformatics/degenotate). Divergence time between pairs of strains was then estimated using the same formula as in (Peter *et al*., 2018) in R. Divergence time per pair of clades was then summarized by the median of all pairwise comparisons between their members. For replicated isolates, a single isolate was randomly selected for inclusion, and isolate Pundurs 1 was excluded from the Baltic group.

### Public datasets used in this study

For Figure 1B, 157 assemblies by (Gallone *et al*., 2016) (BioProject PRJNA323691), 8 sequencing runs from (Preiss *et al*., 2018) as well as genome assemblies of 4 *S. cerevisiae* strains SX1, SX3, HN10, and HN19 (Duan *et al*., 2018) (GCA_003277725.1, GCA_003277085.1, GCA_003275955.1, GCA_003275885.1) were retrieved from NCBI. For variant calling and phylogenies (Figure 2), a panel of raw sequencing data was created according to Supplementary Data 1 using selected strains from the panel compiled by (Tellini *et al*., 2024) adding data from (Preiss *et al*., 2024). A larger phylogeny of 1,269 samples (Supplementary Figure 2) was created from data listed in Supplementary Data 8 based on the same panel. In both panels, clades with names containing (‘mix’, ‘other’, ‘hybrid’, ‘mosaic’, ‘introgressed’) were excluded, except for ‘MosaicsBeer’. Accessions of ‘Chico’ sequences used in Figure 1C are listed in Supplementary Data 9 (Large *et al*., 2020).

### Workflow implementation and reproducibility

All workflows were managed using Snakemake, with Conda environments specified for each step to ensure reproducibility and executed on a high-performance compute server under Linux. All code repositories are publicly available on GitHub. An exhaustive list of code created for the analyses described herein is given in Supplementary Data 6.

## Supplementary Information

Supplementary Data and Information has been deposited in Figshare under DOI: 10.6084/m9.figshare.31709038

## Declarations

### Ethics approval and consent to participate

Ethics approval not applicable. All kveik donors provided informed consent for the use of their strains in research.

### Consent for publication

Not applicable.

### Availability of data and materials

Raw sequencing data and assemblies have been deposited in the European Nucleotide Archive (ENA) under umbrella project PRJEB90469 (sequencing data: PRJEB89706; assemblies: PRJEB90461, PRJEB90466, PRJEB90467, PRJEB90468). Variant matrices have been deposited in Figshare (Dondrup, 2025).

A comprehensive summary of all run accessions assigned is given in Supplementary Data 7. *S. cerevisiae* strains isolated in this study have been preserved in the kveik strain bank (“Kveikbank”) operated jointly by the Western Norway Culture Academy and NIBIO and can be obtained from the authors upon reasonable request.

### Competing interests

The authors declare that they have no competing interests.

### Funding

The work was financed in part by a grant from the Regional Research Council of western Norway, Landbruksdirektoratet, Norway, and in part by research funding from the Western Norway Culture Academy and NIBIO. The Genomics Core Facility (GCF) at the University of Bergen, which is a part of the NorSeq consortium, provided services to the project; GCF is supported in part by major grants from the Research Council of Norway (grant no. 245979/F50). OI and HS were supported by the European Union’s Horizon 2020 research and innovation programme under grant agreement no. 956351 “ChemArch”. Bioinformatics analyses were carried out under the framework of ELIXIR-Norway (https://elixir.no) funded by the Research Council of Norway (Elixir3, grant no.322392). The funders had no role in the conceptualization, design, data collection, analysis, decision to publish, or preparation of the manuscript.

### Author contributions

HGE, AOM and MD: study design. HGE, AOM, AE, DD and MD: sample and data collection. HGE, LKH, AE and JB: laboratory sample analysis. MD, DD, HGE, OI, HS, SNG, TM: bioinformatic analysis, data analysis, and interpretation. HGE, MD, AOM, SH, AE, JB, OI, HS and TM: manuscript writing and editing. All authors contributed to and approved the final version of the manuscript.

## Acknowledgements

We thank the kveik owners for sharing their yeast with us for this study, and Rita Holdhus, Jorunn Børve, Lorena Butinar and Ingunn Øvsthus for advice and assistance. David Kniha is acknowledged for skillful assistance with the map (Figure 1). The authors would like to thank Grigoris Amoutzias and Bram Danneels for helpful discussions on the study and reviewing the draft manuscript.

## Notes

https://doi.org/10.6084/m9.figshare.31709038

### Competing Interest Statement

The authors have declared no competing interest.

### Summary of Updates

- The title of the manuscript has been changed to "Genomic evidence of early-diverging domesticated lineages in Norwegian farmhouse yeast". - The abstract has been changed to a non-structured format. - A new study by Bircham et al. (2026) has been included in the Results and the Discussion section. - We have clarified and harmonized the terminology regarding our samples, isolates, and original brewing cultures. - We have clarified the Materials & Methods section to better represent our grouping strategy and the phylogenetic methods used. - We have added to and clarified the Discussion section.

