## Supplementary Information and Figures for "Genomic evidence of early-diverging domesticated lineages in Norwegian farmhouse yeast"

Supplementary Notes, Figures and Tables

Michael Dondrup^1*^, Atle Ove Martinussen^2^, Lisa Karine Haugland^3^, Jonas Brandenburg^4^, Oya Inanli^5,6^, Abdelhameed Elameen^3^, David Dolan^1^, Sushma Nagaraja Grellscheid^1,9^, Snorre B. Hagen^7^, Hannes Schroeder^5^, Tor Myking^8^, and Hans Geir Eiken^7*^

^1^Department of Informatics, Computational Biology Unit, University of Bergen, Norway, ^2^Western Norway Cultural Academy, Voss, Norway, ^3^Division of Biotechnology and Plant Health, Norwegian Institute of Bioeconomy Research, Ås, Norway, ^4^Michael Sars Centre, University of Bergen, Norway, ^5^Globe Institute, Faculty of Health and Medical Sciences, University of Copenhagen, Denmark, ^6^BioArch, Department of Archaeology, University of York, United Kingdom, ^7^Division of Environment and Natural Resources, Norwegian Institute of Bioeconomy Research, Ås, Norway, ^8^Division of Forestry and Forest Resources, Norwegian Institute of Bioeconomy Research, Ås, Norway, ^9^ Department of Biosciences, Durham University, United Kingdom

(*) Correspondence:

Hans Geir Eiken

Michael Dondrup

### Supplementary Figures

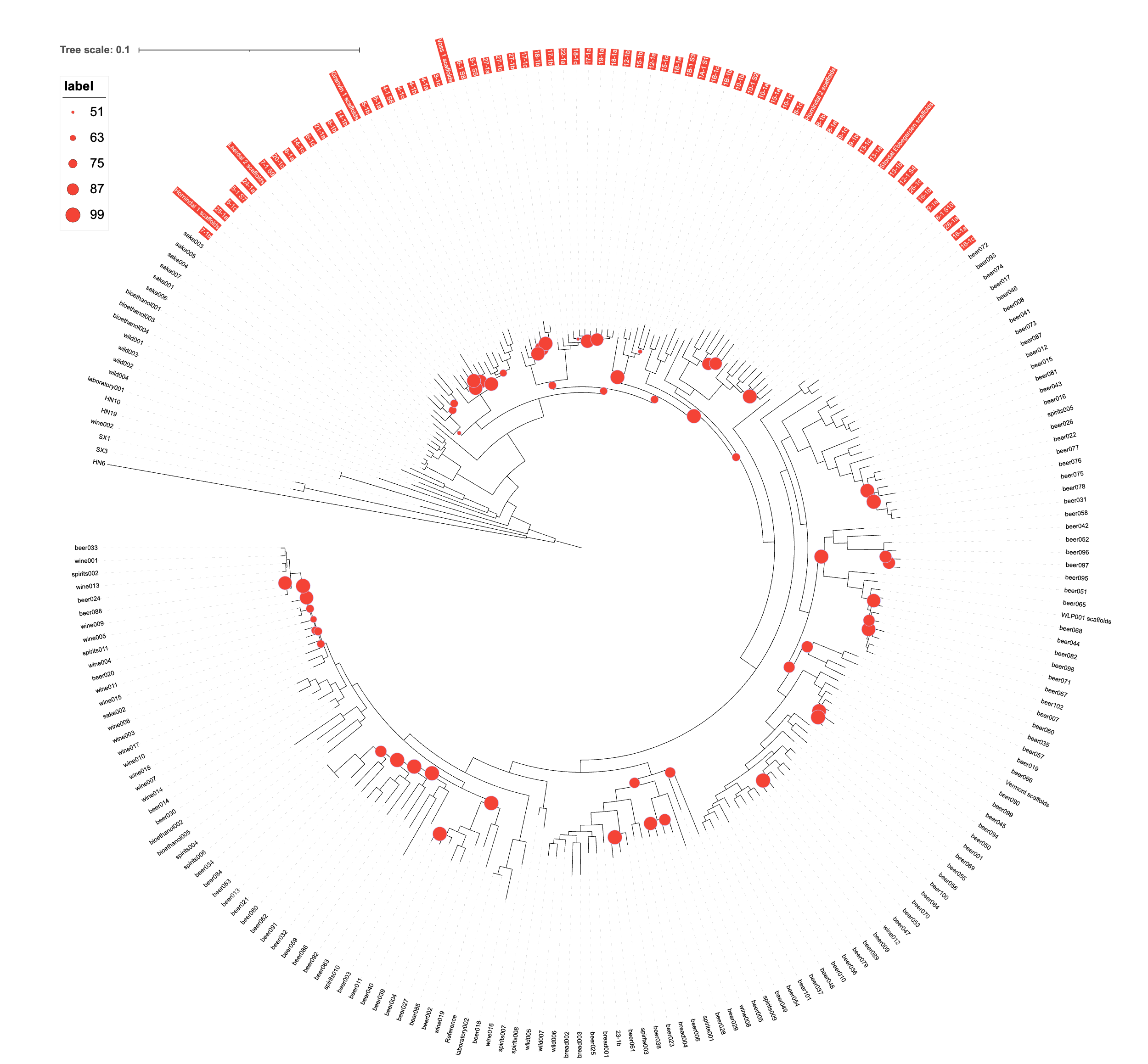

Supplementary Figure 1. Complete whole genome maximum-likelihood (ML) phylogenetic tree of 235 assemblies based on 287,696 parsimony-informative sites as depicted in Figure 1B. In addition to the sequences generated in this study (Supplementary Data 2), we included all strains from the studies of Gallone et al. (2016) and Preiss et al. (2018), as well as additional wild strains from primeval Chinese forests (Duan et al. 2018) to root the tree. Taxon labels of kveik strains are highlighted in red. Bootstrap percentages are given by circles of proportional size for nodes with support < 100%. Substitution model chosen according to BIC: TVM+F+ASC+R4

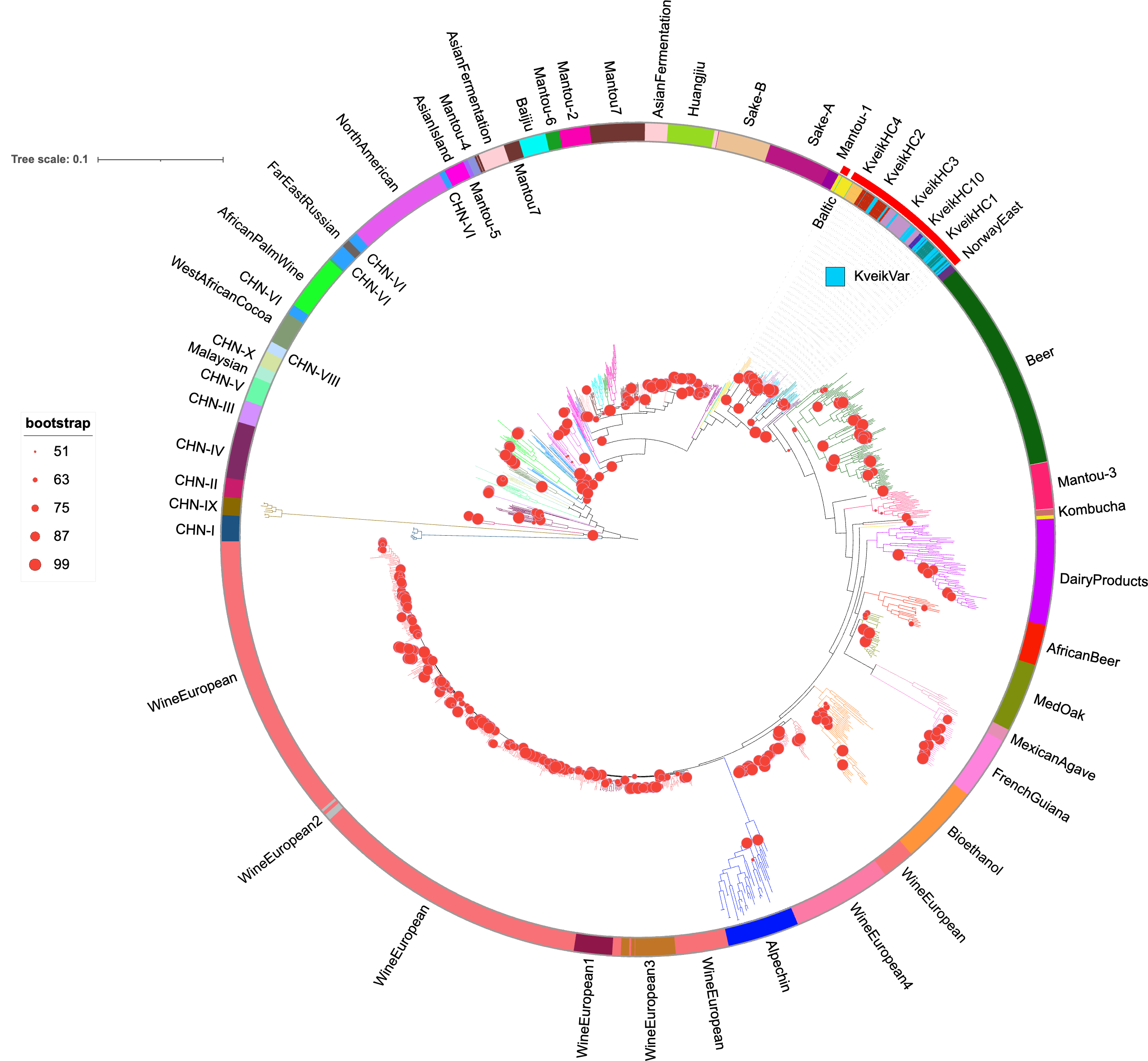

B

A

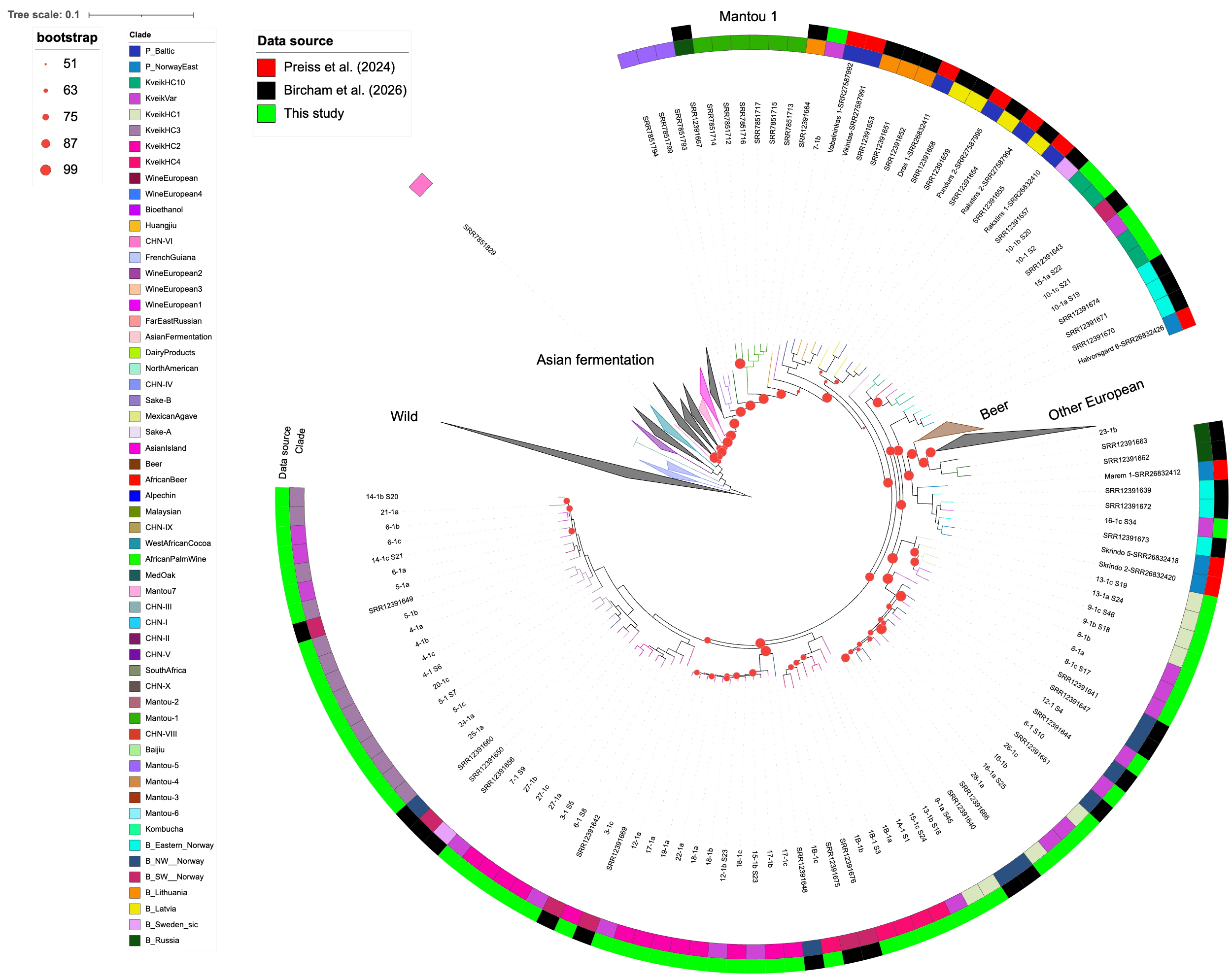

Supplementary Figure 2. A) ML-phylogeny of 1,269 S. cerevisiae isolates including kveik using the panel compiled by Tellini et al. (2024). The tree was based on 567,383 parsimony-informative sites. Major clades have been annotated. B) ML-phylogeny of 1,306 isolates also including data from the recent study of Bircham et al. (2026) for comparison, based on 568,374 parsimony-informative sites, most clades except farmhouse cultures were collapsed. Data source is indicated by color (outer ring), clade assignments of taxa are indicated by color (inner ring). Both trees were computed using the GTR+ASC+G substitution model in IQ-Tree with 1,000 ultra-fast bootstrap iterations. Only bootstrap support values < 100% are annotated with red dots with proportional diameter. Both trees have been rooted on the joint CHN-I, CHN-IX clade. Taxon information for the trees is provided in Supplementary Data 8. The trees are available for interactive browsing via iTOL A) <https://itol.embl.de/tree/9222021523975201760864481> and B) <https://itol.embl.de/tree/92220215239253281780832119>

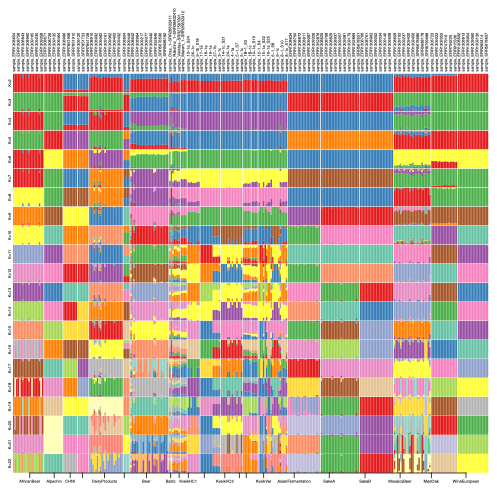

Supplementary Figure 3. Full ADMIXTURE bar-graph over all K from 2 to 22. Minimal cross-validation error is achieved at K=22. This graph also includes the KveikVar group containing isolates that did not cluster consistently and showed variability across sequencing of repeated colonies from the same culture.

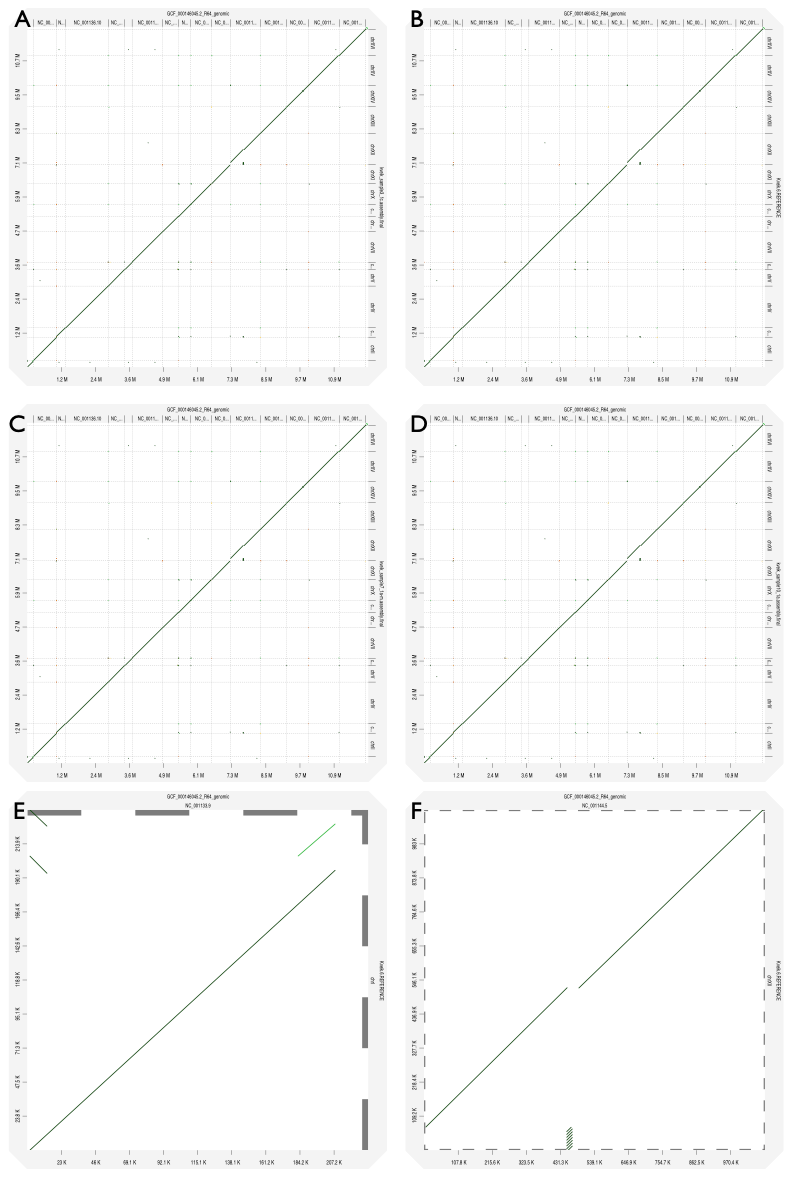

Supplementary Figure 4. Dot plots of the full collapsed long-read kveik assemblies aligned against the R64 reference genome. Assemblies: A) sample 3_1c, B) sample 6_1a, C) sample 7_1a, D) sample 10_1a. E, F) Chromosomal dot plots with the largest structural variations between kveik sample 6 and the reference for E) chrI and F) chrXII.

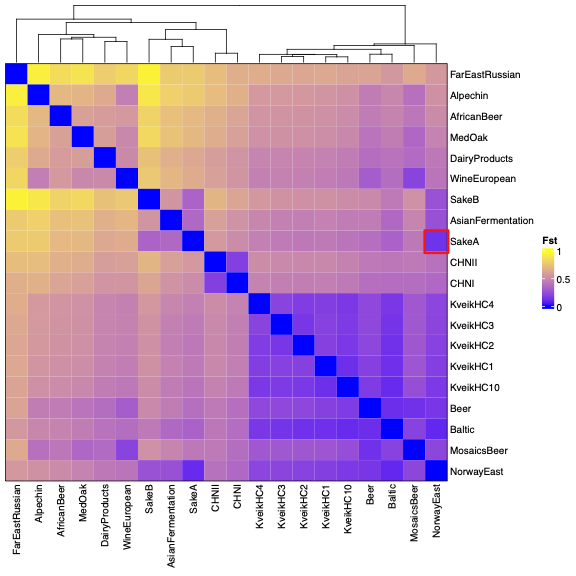
Supplementary Figure 4. Clustered heatmap of genetic distances (as pairwise F_ST_ values) between groups of the variant panel. An outlier with low genetic distance between the eastern Norwegian group and SakeA has been highlighted in red.

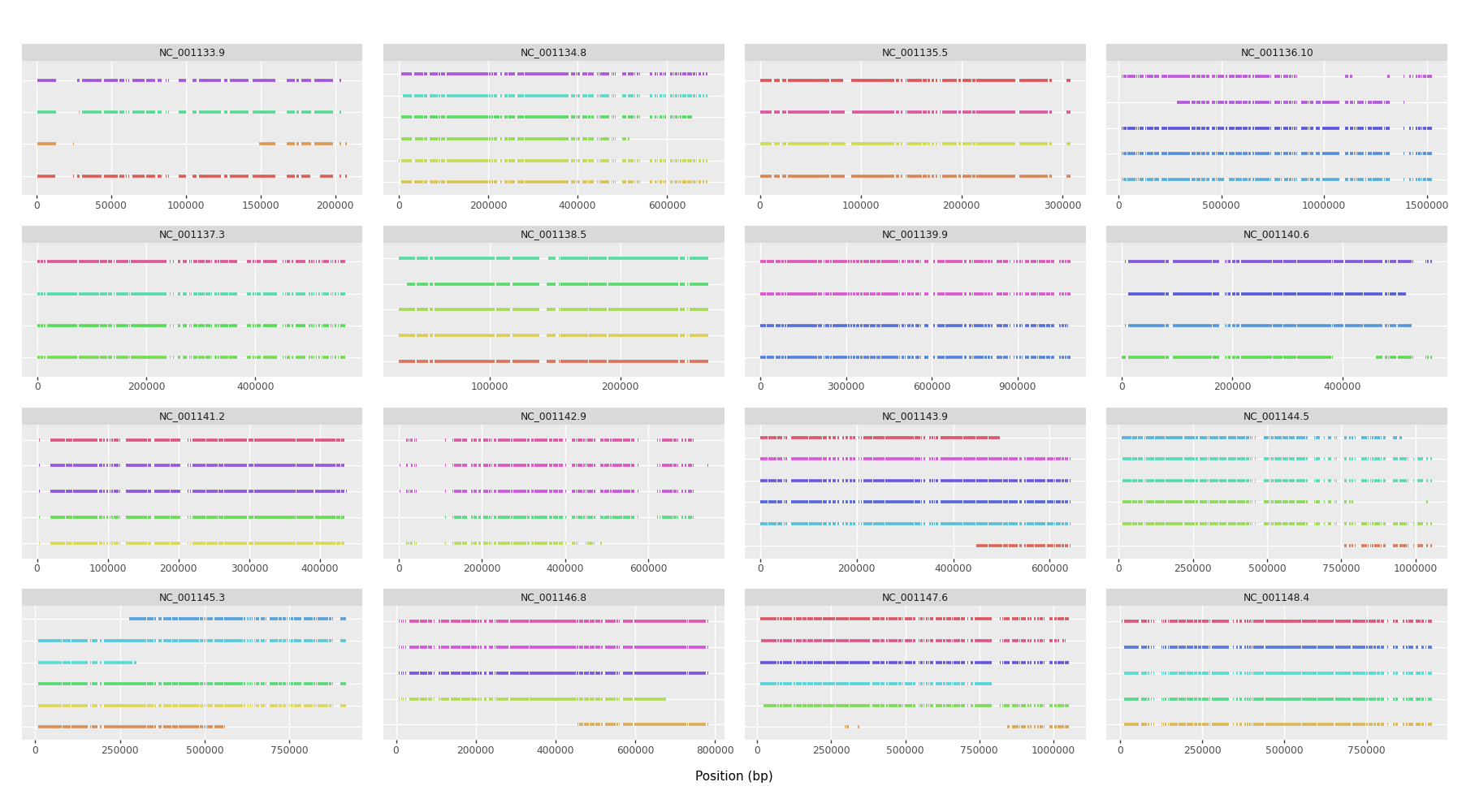

Supplementary Figure 5. Cleaned phased sequence clusters by NPhase aligned to nuclear chromosomes of the S288_R64 reference assembly for sample 3_1c.

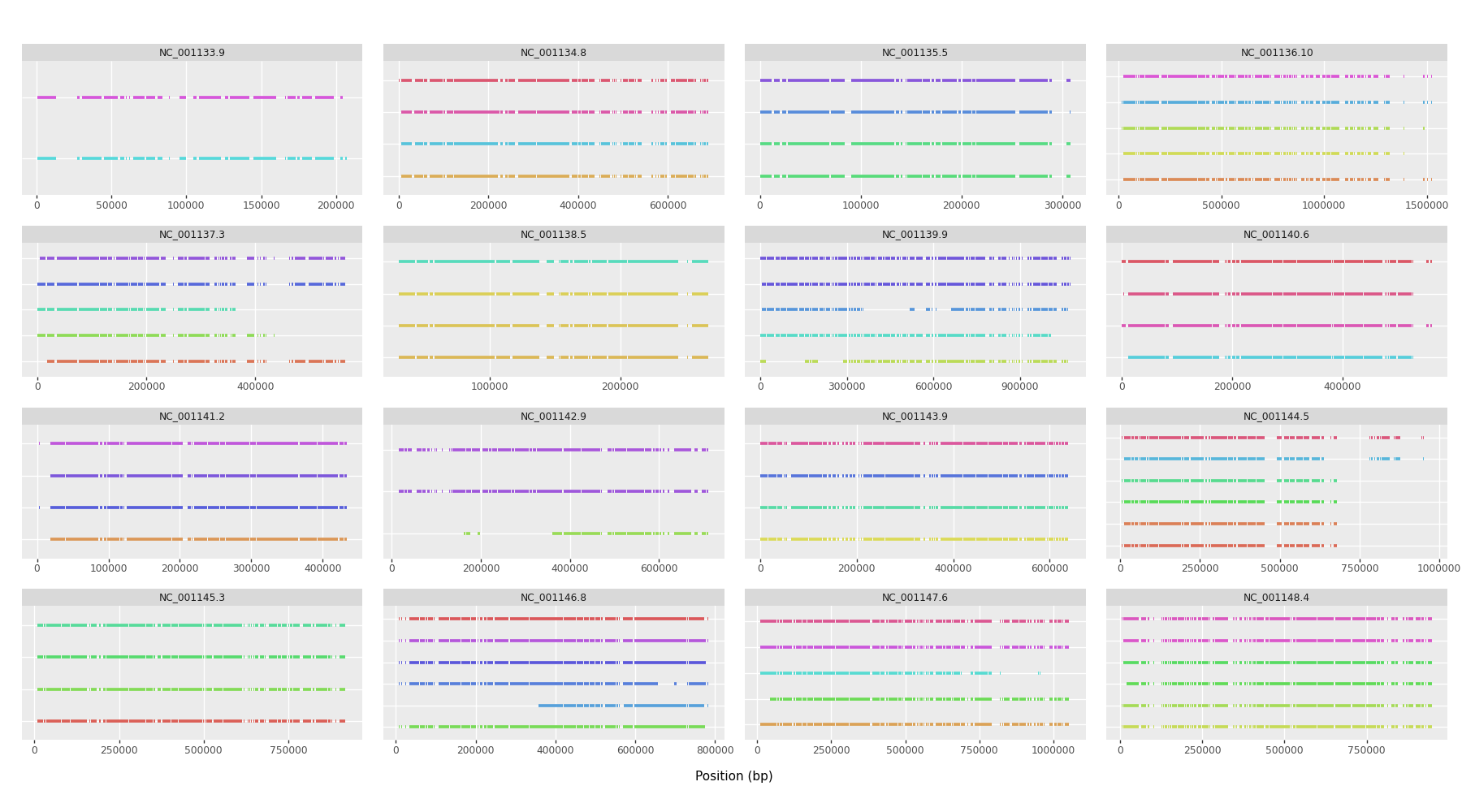

Supplementary Figure 6. Same as above for sample 6_1a.

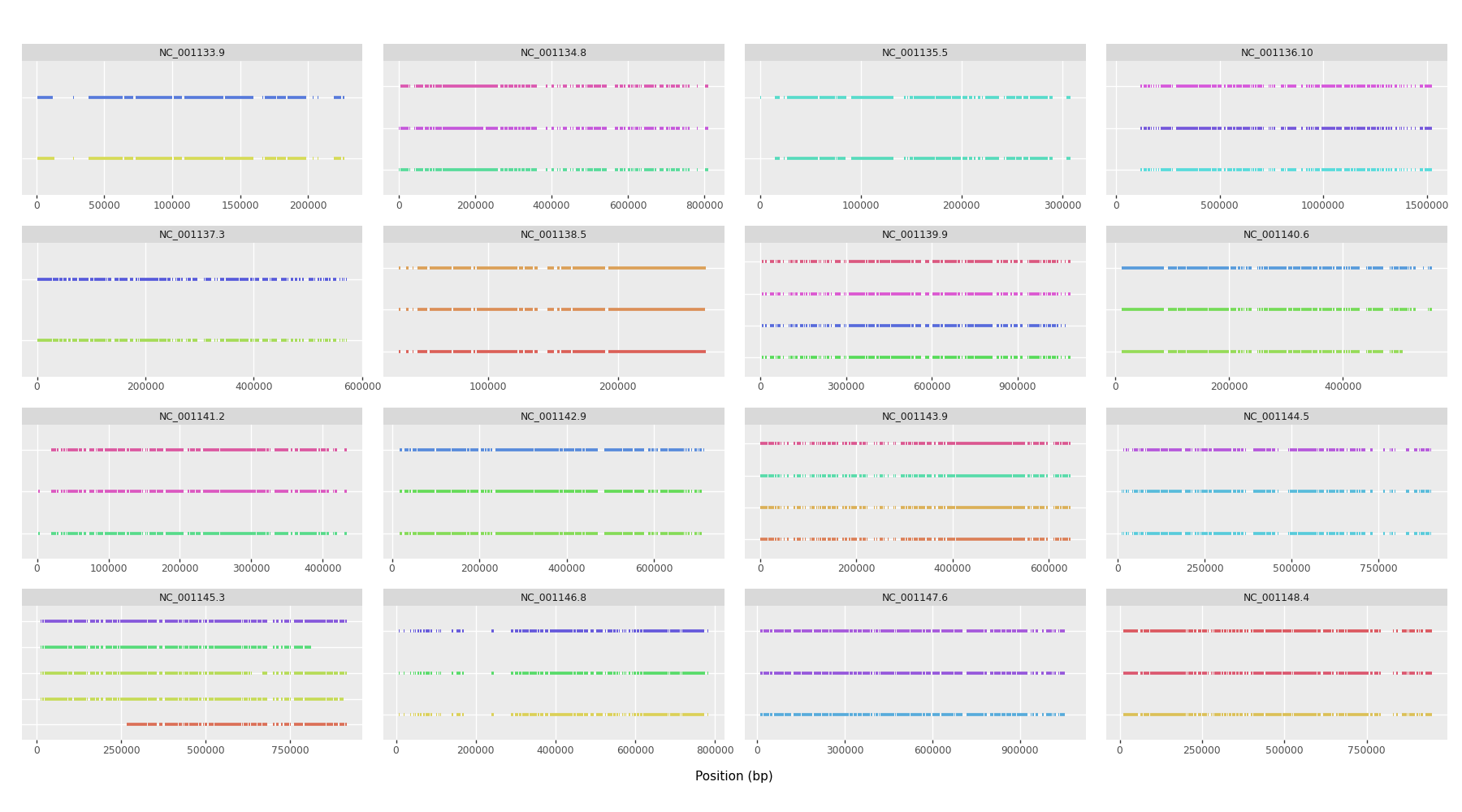

Supplementary Figure 7. Same as above for sample 7_1a.

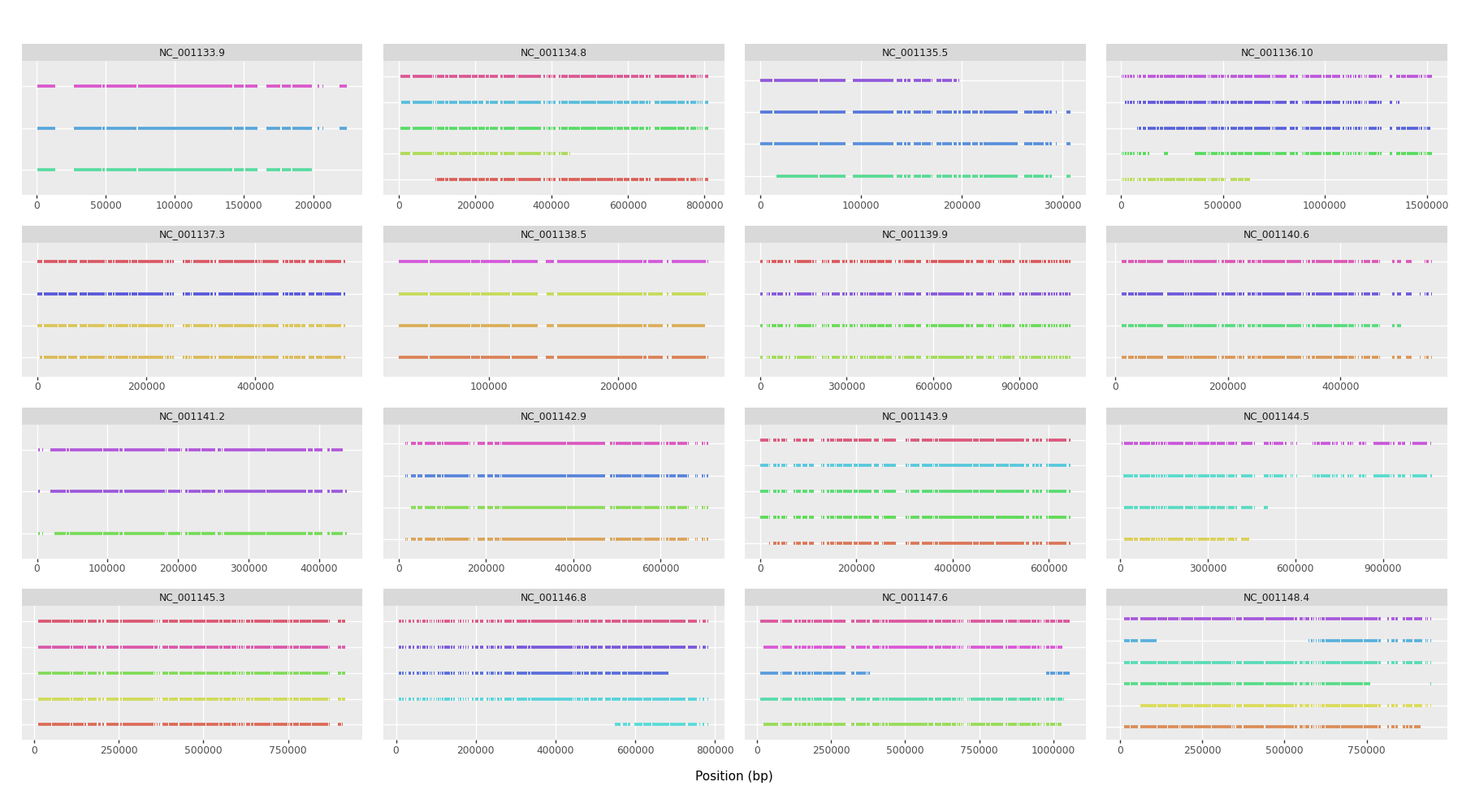

Supplementary Figure 8. Same as above for sample 10_1a.

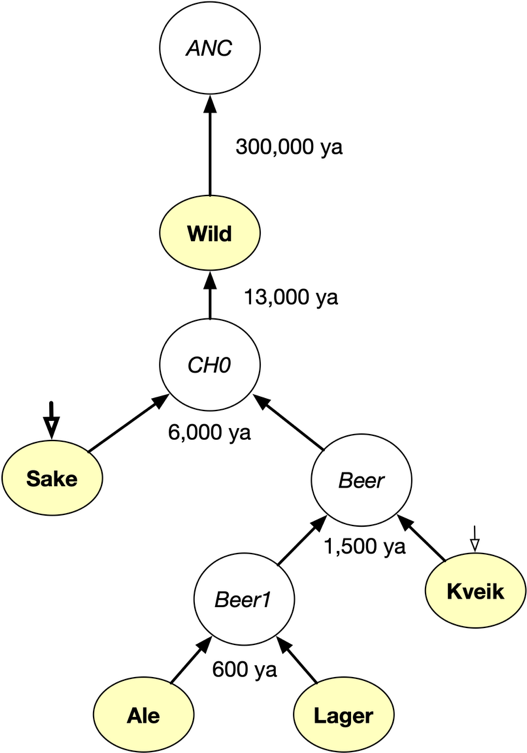

Supplementary Figure 9. Simplified demographic model of S. cerevisiae evolution common to all settings used to simulate genomic data under a coalescent model with msprime. Times are given in years ago (ya) for each population split. Names of sampled populations are given in bold face. In some models, single or multiple instantaneous bottlenecks of variable strength and at different points in time were added to some populations (Sake, Kveik, indicated by open arrowheads). Exact parameters for each model are found in the source code of the model control files.

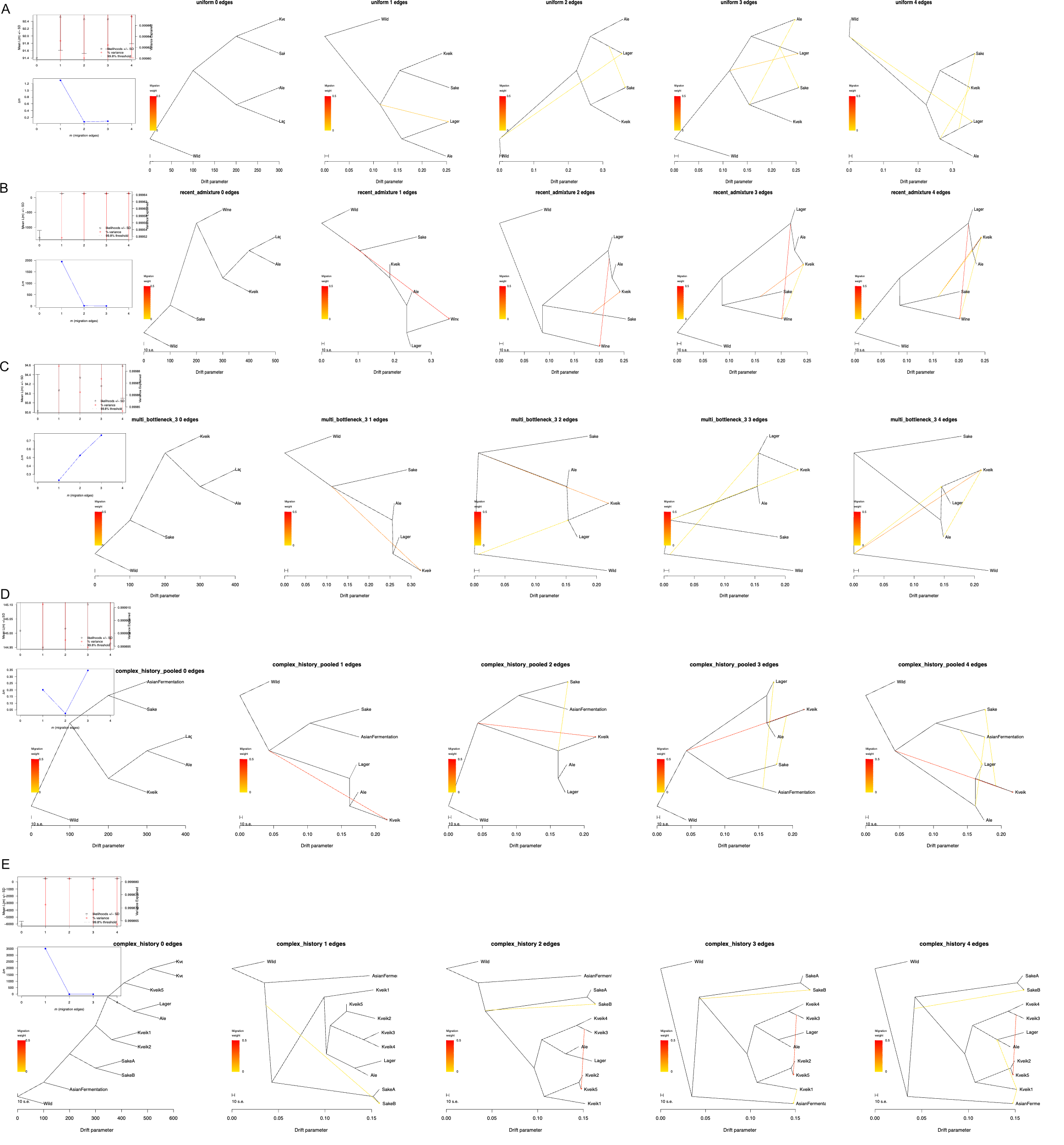

Supplementary Figure 10. TreeMix output generated from variants simulated with msprime under coalescent theory using different demographic models. OptM plots, commonly used to evaluate the optimal number of migration edges, are given in the leftmost column. Tree plots are given for 0 to 4 migration edges in each row. A) Variants were generated from a model with no bottlenecks or admixture and uniform number of samples per group. B) Demographic model including admixture from “Sake” and “beer” into “Kveik”. An additional “Wine” group was added to the model. C) Demographic model with a single strong instantaneous bottlenecks of high strength added to “Sake” and multiple weaker and earlier bottlenecks added to “Kveik”. No admixture was added to any population in this model. D, E) A more complex model of S. cerevisiae was used to demonstrate effects of pooling groups with complex population structure on TreeMix analyses. The model contains five ‘Kveik’ groups with various divergence times and bottlenecks of low strength, a mixed group ‘Kveik5’ - like KveikVar - as well as two ‘Sake’ groups with strong bottlenecks, an ‘AsianFermentation’ group, as well as shadow migration. Both analyses are identical, except for the way samples were annotated. In D) samples from five simulated ‘Kveik’ groups and two ‘Sake’ groups were pooled into collapsed groups, while in E) ‘Kveik1’ – ‘Kveik5’ and ‘SakeA’ and ‘SakeB’ are treated as separate clusters. A ‘phantom migration’ signal into the collapsed ‘Kveik’ group is visible in D) while in E) a migration signal from ‘Kveik3’ into ‘Kveik5’ is observed.

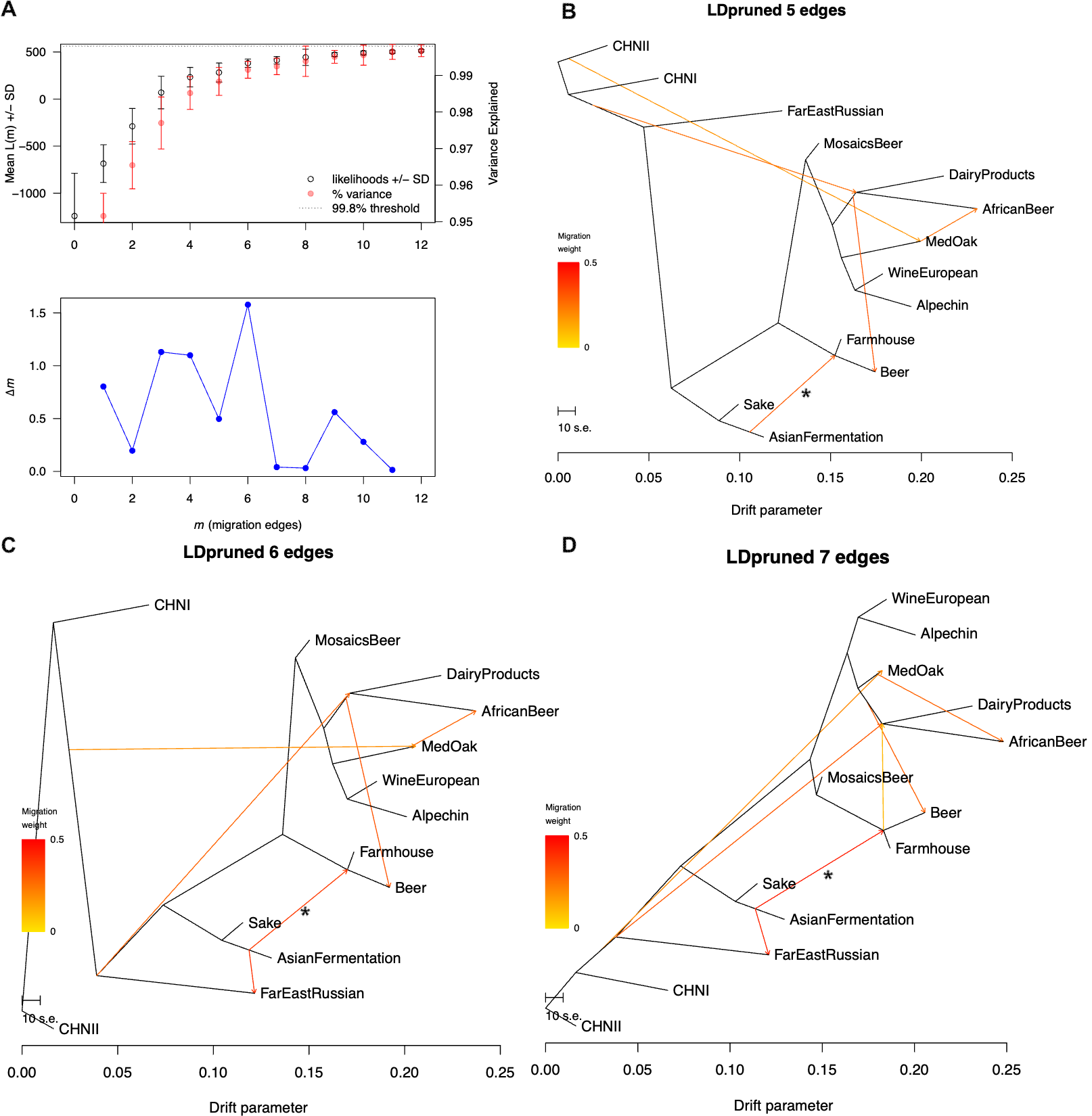

Supplementary Figure 11. Outcome of TreeMix analysis when modeling a structured group as a single panmictic “Farmhouse” population. TreeMix analysis was run on the same data and using the same methods as described in the Methods section. However, Norway East, Baltic, and kveik strains were all grouped into a single hypothetical “Farmhouse” group, and SakeA and SakeB were grouped into “Sake” similar to the approach taken by Preiss et al. 2024. OptM plot (A) suggests 6 as optimal number of migrations. TreeMix graphs with 5 to 7 edges (B-D) show a strong -likely phantom-migration- weight (*) from Sake/AsianFermentation into the Farmhouse/Beer group.

### Supplementary Tables

Supplementary Table 1. Counts of genomic feature types (with respect to the S288c_R64 reference genome) affected by at least one structural variant in all four strains sequenced by Oxford Nanopore Technology (see Supplementary Data 3).

| Feature type | Frequency |
| --- | --- |
| CDS | 258 |
| exon | 220 |
| gene | 217 |
| mRNA | 201 |
| long_terminal_repeat | 178 |
| mobile_genetic_element | 48 |
| origin_of_replication | 18 |
| telomere | 17 |
| tRNA | 10 |
| ncRNA | 2 |
| RNase_MRP_RNA | 1 |
| rRNA | 1 |
| snoRNA | 1 |
| snRNA | 1 |

### Supplementary Notes

#### Traditional Farmhouse Brewing with Kveik

The following detailed protocol outlines the farmhouse kveik brewing process. The description is based on notes from a brewing workshop at a local smokehouse (“Eldhus”) with tradition bearer Sigmund Gjernes in November 2022.

**1. Preparation of juniper infusion:** Fresh juniper branches are placed in the bottom of a copper kettle. Water is added and heated on an open fire to approximately 82°C for 1-1.5 hours. The water is then passed through a sieve and left to cool to 68°C.

**2.** **Mashing:** Malted barley is combined with the juniper-infused water. The mixture is maintained at around 68°C with constant mixing and covered for 4 hours to allow the enzymatic conversion of starch into sugars. This step also kills most, or all, microorganisms present in the malted barley, juniper, and tank.

**3. Lautering:** The mash is drained slowly, and liquid wort passes through the juniper branches at the bottom of the tank, which serve as a natural filter.

**4. Boiling:** The wort is transferred to a copper kettle and boiled for approximately four hours. During this process, the wort is exposed to remove juniper oils and proteins. Again, bacteria and fungi are killed.

**5. Cooling:** After boiling, the wort is cooled to approximately 40°C.

**6.** **Yeast** **pitching** **and** **fermentation:** In farmhouse brewing, liquid kveik yeast, traditionally passed down through generations, is added to the 40°C wort. This is followed by a practice of “shouting” or “yelling" to the kettle, which is believed to enhance fermentation. Fermentation is rapid and typically completed within two days.

**7.** **Conditioning** **and** **storage:** Beer is often consumed fresh, though it may be briefly conditioned to allow the flavors to mature.

#### Admixture Simulation

##### Methods

The workflow was designed to evaluate TreeMix with simulated data using different demographic models, both with and without admixture. msprime (version 1.3.3) (Baumdicker et al. 2022) was used to simulate genetic data for various yeast populations under a coalescent model incorporating different demographics such as uniform, recent admixture, multi-bottleneck, and bottlenecks with different group sizes. Parameters for msprime included a fixed effective population size (isize = 1e6), generations per year (2900), and mutation rate (rate = 1.67e-10/generation). The simulations were performed with a sequence length of 12 million base pairs and random seeds for reproducibility. A random sample of 20,000 SNPs was taken from the simulated files for each model except for Models D and E (Supplementary Figure 10) where 100,000 SNPs were used. TreeMix was run allowing for 0 to 4 migration events with 100 bootstrap iterations per dataset. The derived trees were then assessed with OptM. The workflow was implemented in Snakemake, the source code is available on Github (<https://github.com/mdondrup/admixture_simulation>).

All demographic models included five simulated *S. cerevisiae* populations from which samples were generated: “Kveik”, “Lager”, “Ale”, “Sake”, and “Wild”. Four hypothetical ancestral populations were also defined: “ANC” (the ancestral equilibrium population), “CH0” (the original *S. cerevisiae* “out-of-China” population), “Beer” (the ancestral population of all Beer and Kveik), and “Beer1” (the ancestor of “Ale” and “Lager”). Modern sake strains are thought to have passed through a strong and recent population bottleneck (Ohya and Kashima 2019). Therefore, in some of our models, instantaneous bottlenecks of different strengths were added to the simulated “Sake” and “Kveik” groups. In the “Kveik” group, multiple -but weaker- consecutive bottlenecks are intended to resemble the internal collapsed population structure inferred from ADMIXTURE analysis. Demographic parameters common to all models are depicted in Supplementary Figure 1. The model “recent_admixture.py” also contains an added admixture signal while none of the other models contains admixture. In the models “complex_history.py” and “complex_history_pooled.py”, additional structure and subgroups of Kveik and Asian fermentation strains as well as some different initial population sizes were introduced and a recent admixture pulse kveik subgroups (Kveik2 + Kveik3 → Kveik5) between, but not across major lineages, were added. Both complex models differ only in how they treat subgroups of “Kveik” and “Sake” either as one pooled group or as individual groups in the subsequent TreeMix analysis.

##### Results

Despite the simplified demographic models underlying our simulation experiments, with constant population size and no other structure than pulse bottlenecks, various outcomes contained pronounced false-positive migration signals. Uniform models did only produce very weak migration signals (Supplementary Figure 10A). Notably, models containing multiple bottlenecks of different strength in the simulated “Kveik” and “Sake” populations resulted in strong spurious migration signals into the “Kveik” group (Supplementary Figure 10C), with the strength of the signal apparently depending on the difference between the bottleneck events. These signals resemble the outcome when admixture is added to the model, even though neither source nor targets of admixture signal were correctly identified (Supplementary Figure 2B). In addition, Δm values calculated by OptM, which is commonly used to assess the optimal number of migration events point at that increasing an increasing number of migration edges may yield better results (Supplementary Figure 10C).

To investigate the possible effects of pooling, a more complex and realistic model was developed, including a highly structured simulated ‘Kveik’ group that was analyzed once by pooling all samples in the ‘Kveik’ group, and once by treating them separately (Supplementary Figure 10D&E). The effect of pooling samples was comparable to the multiple bottleneck setting, while modeling populations directly resulted in a more accurate resolution of the demographic history, including the resolution of the modelled admixture into ‘Kveik5’ (Supplementary Figure 10E).

This confirmed our initial hypothesis that “phantom-migration” signals in TreeMix are readily evoked by unmodeled population structure and therefore migration signals into e.g. a collapsed Kveik group need to be interpreted with utmost caution.
